# Cooperation between C*aenorhabditis elegans* COMPASS and condensin in germline chromatin organization

**DOI:** 10.1101/2020.05.26.115931

**Authors:** M. Herbette, V. Robert, A. Bailly, L. Gely, R. Feil, D. Llères, F. Palladino

## Abstract

Deposition of histone H3 lysine 4 (H3K4) methylation at promoters by SET1/COMPASS is associated with context-dependent effects on gene expression and local changes in chromatin organization. Whether SET1/COMPASS also contributes to higher-order chromosome structure has not been investigated. Here, we address this question by quantitative FRET (Förster resonance energy transfer)-based fluorescence lifetime imaging microscopy (FLIM) on *C. elegans* germ cells expressing histones H2B-eGFP and H2B-mCherry. We find that SET1/COMPASS subunits strongly influence meiotic chromosome organization, with marked effects on the close proximity between nucleosomes. We further show that inactivation of SET-2, the *C. elegans* homologue of SET1, or CFP-1, the chromatin targeting subunit of COMPASS, strongly enhance chromosome organization defects and loss of fertility resulting from depletion of condensin-II. Defects in chromosome morphology resulting from conditional inactivation of topoisomerase II, another structural component of chromosomes, were also aggravated in the absence of SET-2. Combined, our *in vivo* findings suggest a model in which the SET1/COMPASS histone methyltransferase complex plays a role in shaping meiotic chromosome in cooperation with the non-histone proteins condensin-II and topoisomerase.

## Background

In different species, from yeast to mammals, chromatin modifying complexes and histone post-translational modifications (PTMs) contribute to higher-order chromatin structure (Rowley & Corces, 2018). While the spatial configuration of chromatin is essential to ensure fundamental processes from gene expression to cell divisions, how higher-order structures are formed in various cellular processes remains unclear.

Mitosis and meiosis are essential cellular processes that require restructuring and reorganization of chromatin architecture, and are both associated with specific changes in histone modifications. During mitosis, extensive compaction of chromatin is associated with histone H3 serine 10 phosphorylation (H3S10ph), H4 lysine-20 mono-methylation (H4K20me1), and a dramatic reduction in overall histone acetylation (Antonin & Neumann, 2016; Zhiteneva *et al*, 2017; Beck *et al*, 2012). Specific histone PTMs, including H3 lysine-4 tri-methylation (H3K4me3), are also associated with meiotic double strand breaks (DSBs) during recombination, and are dynamically altered during meiotic progression (Borde *et al*, 2009; Buard *et al.*, 2009). In addition, during mammalian spermatogenesis, a large fraction of the observed dynamic changes in H3K4me3 do not coincide with either gene promoters, or double strand breaks (DSB) (Lam *et al*, 2019), suggesting additional functions that remain to be explored.

SET1 family histone methyltransferases act in large multi-subunit complexes known as COMPASS (Complex Proteins Associated with Set1) (Eissenberg & Shilatifard, 2010; Dehe *et al,* 2006) to deposit H3K4me3 at promoters of actively transcribed genes. At promoters, levels of COMPASS-dependent H3K4 methylation generally correlate with transcription levels, but evidence for an instructive role for H3K4me3 in transcription is lacking, and recent data suggest that its role depends on different chromatin and cellular contexts (Howe *et al*, 2017).

Studies of individual components of COMPASS are consistent with functions in various aspects of chromatin organization. For example, binding of yeast COMPASS component Spp1/CFP1 to H3K4 is required to tether loop formation and DNA cleavage at sites of double strand breaks (DSBs) (Borde *et al*, 2009; Acquaviva *et al*, 2013; Sommermeyer *et al*, 2013). Additional H3K4-dependent and -independent roles for yeast Set1 in genome organization through long-range clustering of retrotransposon loci have also been described (Mikheyeva *et al*, 2014; Lorenz *et al*, 2012). Inactivation of CFP1 in developing mouse oocytes results in defects in meiotic oocyte maturation, spindle assembly and chromosome alignment, with only minor effects on transcription (Sha *et al*, 2018). (Yu *et al*, 2017).

Inactivation of *set-2*, the single SET1 homolog in *C. elegans*, results in defective patterns of H3K4me3 in the germline, progressive misregulation of the germline transcriptome, increased genome instability, and loss of germ cell identity leading to sterility (Herbette *et al*, 2017; Li & Kelly, 2011; Robert *et al*, 2014; Xiao *et al*, 2011). Whether these defects reflect a direct role of SET-2 in transcription, or a more general role in germline chromatin organization is not known. We previously found no correlation between COMPASS-dependent H3K4me3 at promoters and transcription in *C. elegans* embryos, which is consistent with observations in other organisms (Clouaire *et al*, 2012, 2014; Beurton *et al*, 2019; Lenstra *et al*, 2011; Weiner *et al*, 2012) and argues against a direct role in transcription. Likewise, increased genome instability in *set-2* mutant germlines was not associated with defects in the induction of the DNA damage response (DDR) pathway, consistent with downstream effects in the DNA repair process (Herbette *et al*, 2017). Together, these recent observations suggest that SET-2 may play roles in modifying chromatin structure independently of specific changes in gene expression.

In this study, we used fluorescence lifetime imaging microscopy (FLIM) for Förster Resonance Energy Transfer (FRET) measurements to directly assess changes in chromatin compaction in live animals lacking COMPASS components. We find that FRET between fluorophore-tagged nucleosomes is dramatically decreased in meiotic cells from *set-2* and *cfp-1* mutant animals, consistent with a structural role for COMPASS in germline cells affecting close nucleosome proximity in the nucleus. Loss of either *set-2* or *cfp-1* enhanced chromosome organization defects resulting from depletion of condensin-II, a major regulator of chromosome structure (Hirano, 2012), again consistent with a structural role for COMPASS in chromosome organization. Cooperation between SET-2 and condensin-II in the germline was independently confirmed in animals carrying a conditional allele of condensin-II subunit *hcp-6*. We further show that *set-2* inactivation aggravates the germline phenotypes resulting from conditional inactivation of topoisomerase II, another major structural component of chromosomes (Adachi *et al*, 1991; Hirano & Mitchison, 1993; Uemura *et al*, 1987). Altogether, our data indicate that COMPASS contributes to the organization of germline chromosome architecture in cooperation with condensin and topoisomerase-II, and evoke the possibility that COMPASS-related complexes and H3K4 methylation are broadly involved in higher-order chromatin structure.

## Results

### Nanoscale chromatin compaction is decreased in *set-2* mutant germlines

In the *C. elegans* germline, meiotic nuclei are arranged in a temporal-spatial order, with the distal end of the gonad containing mitotically proliferating nuclei. Homolog pairing initiates downstream in the “transition zone”, followed by the pachytene stage, during which synapsed chromosomes appear in DAPI-stained nuclei as discrete, parallel tracks. More proximally, nuclei exit pachytene, enter diplotene, and cellularized oocytes containing condensed homologs are formed (Kimble & Crittenden, 2005).

H3K4me3 is detected on chromatin in all germline nuclei, from the distal mitotic region through the meiotic stages and into diakinesis (Figure S1A, (Li & Kelly, 2011; Xiao *et al*, 2011). In germlines from animals carrying the *set-2(bn129)* loss-of-function allele (Xiao *et al*, 2011), H3K4me3 strongly decreases in the distal mitotic region through early-mid pachytene. Levels of H3K4me3 are not visibly altered in late pachytene and diakinetic nuclei of mutant animals, most likely reflecting the additional activity of SET-16/MLL, the only other SET1 family member in *C. elegans* (Figure S1A, (Li & Kelly, 2011; Xiao *et al*, 2011; Fisher *et al* 2010). While reduced H3K4 methylation in *set-2(bn129)* animals at the permissive temperature does not result in obvious fertility defects, a progressive decrease in fertility is observed at the stressful temperature of 25°C, resulting in sterility at the F4-F5 generation (Li & Kelly, 2011; Xiao *et al*, 2011). These observations suggest that H3K4 methylation, or SET-2 itself, play an important role in germline maintenance.

DAPI staining of chromatin revealed no apparent defects in either germline organization, or chromosome morphology in *set-2* mutant animals raised under standard conditions (20°C), or late generation fertile animals at 25°C (data not shown and (Xiao *et al*, 2011). Interestingly however, in late generation (F4) germline nuclei from animals approaching sterility at 25°C, meiotic progression and chromatin compaction were altered, with a loss of the distinctive pachytene nuclei morphology (Figure S1B). Therefore the stressful temperature reveals a role for SET-2 not only in the meiotic process, but also in global genome organization.

Chromatin modifying complexes and histone modifications define the different functional states of chromatin, which in turn contribute to higher-order chromatin organization (Adriaens *et al,* 2018). To investigate how COMPASS influences chromatin architecture specifically in germline cells, we used a recently developed FLIM-FRET technique to quantify changes in chromatin compaction in live animals at the nucleosomal level. The assay is based on the measurement of FRET interactions between fluorescently-labelled core histone GFP::H2B (donor) and mCherry::H2B (acceptor) (Llères *et al,* 2009) co-expressed in the germline from a single transcription unit driven by the germline-specific *pmex-5* promoter to achieve appropriate expression levels (Llères *et al,* 2017). Transgenic animals expressing the two H2B fusion proteins did not show any obvious cell division, growth or reproductive defects (Llères *et al,* 2017). Important features of this system include the following (Llères *et al,* 2017): 1) H2B fusion proteins only represent approximately 4% of total histone H2B, so that only a minute fraction of total nucleosomes is expected to contain both tagged H2B histones as a potential source of intra-nucleosomal FRET; 2) GFP and mCherry are fused to the N-terminus of histone H2B, and the distance separating them from the histone proteins is too large (130 Å on average) to produce significant intra-nucleosomal FRET; 3) FRET occurs efficiently only when the donor and acceptor fluorescent fusion proteins are closely positioned (<10nm) in the 3D nuclear space following chromatin compaction. Importantly, FLIM-FRET also provides accurate quantification due to the independence of the fluorescence lifetime from the relative concentrations of the interacting proteins, and is independent of their diffusion rates (Llères *et al,* 2007). In conclusion, our assay measures close contacts in the nuclear space between distant nucleosomes, and thus provides a read-out of nanoscale chromatin compaction (Lou *et al,* 2019; Sobecki *et al,* 2016; Baarlink *et al,* 2017; Wang *et al,* 2019a).

To carry out the FLIM-FRET assay, we first generated wildtype and *set-2* mutant strains that stably co-express both GFP-H2B and mCherry-H2B fusion proteins (FPs, collectively named “H2B-2FPs” hereafter) from a single transcription unit driven by the germline-specific *pmex-5* promoter (Figure 1A). We confirmed that there was no alteration in the expression of fluorophore-tagged H2B histones in *set-2* mutants (Figure S2A), and fluorescence recovery after photo-bleaching (FRAP) showed that the tagged histones H2B were homogeneously incorporated into chromatin (Figure S2B).

**Figure 1.**
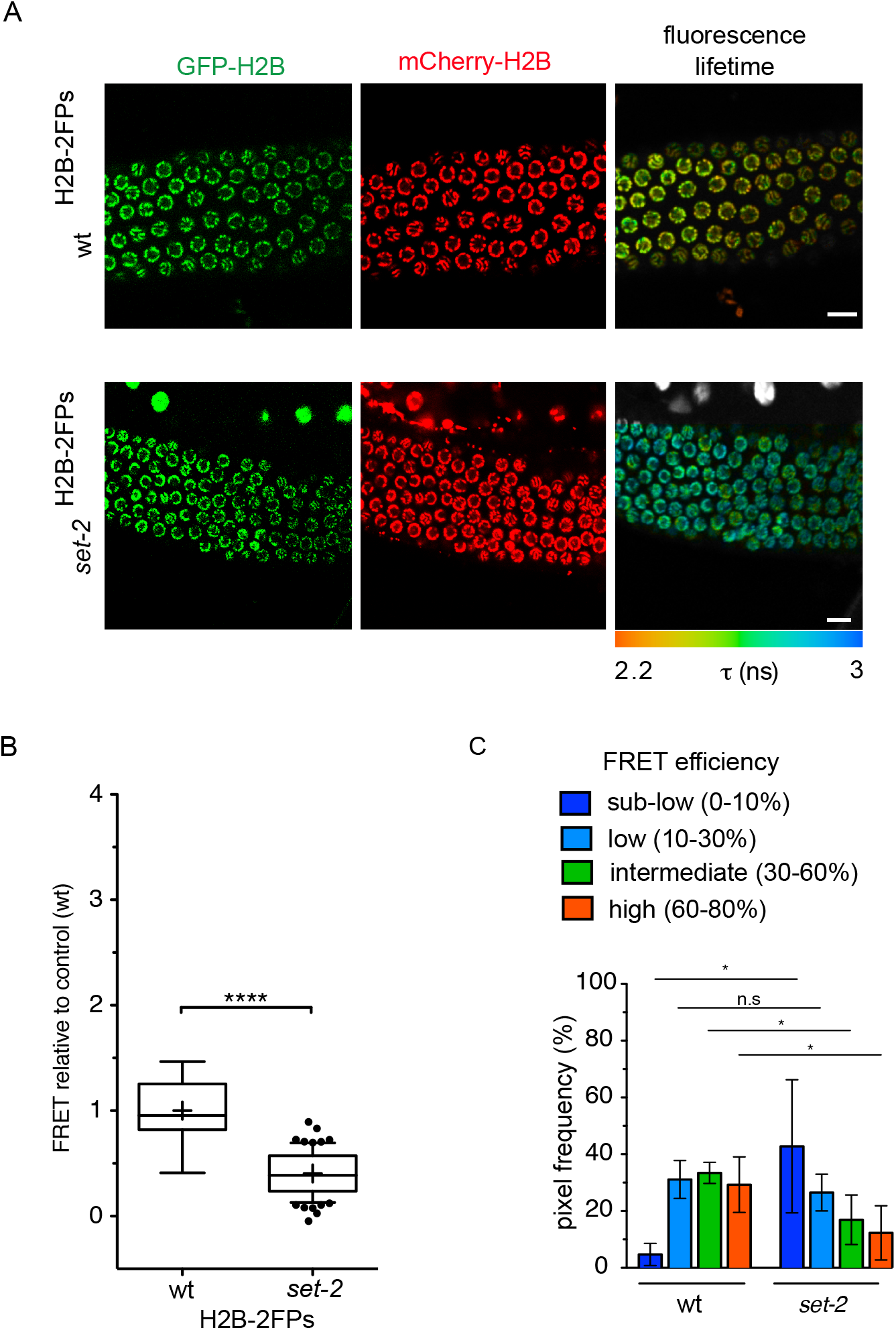
*set-2* inactivation influences nanoscale chromatin compaction in the germline. (**A**) Fluorescence intensities of GFP-H2B (green) and mCherry-H2B (red) from pachytene-stage germ cells expressing H2B-2FPs. FLIM (right) images of H2B-2FPs pachytene-stage cells. The spatial distribution of the mean fluorescence lifetime (τ) at each pixel is shown for wildtype (wt_H2B-2FPs_ cells) (top) or *set-2(bn129)* H2B-2FPs mutant animals (bottom). Fluorescence lifetime values ranging between 2-3 ns are represented using a continuous pseudo-color scale. Scale bars, 10 μm. (**B**) Statistical analysis of the FRET efficiency relative to control (wt), presented as box-and-whisker plots. The mean FRET value is indicated by a cross in each box. ****p < 0.0001 (two-tailed unpaired t test). (**C**) Relative fraction of FRET populations (sub-low, low, intermediate, and high) representing different levels of compaction as defined previously (Llères *et al*, 2017) from wt and *set-2(bn129)* pachytene nuclei. *p < 0.05; (twotailed unpaired t test); n.s, non-significant. n=5 gonads (approx. 350 nuclei) for wt, n=6 gonads (approx. 430 nuclei) for *set-2(bn129).*

Comparative FLIM-FRET analysis of *set-2(bn129)*_H2B-2FPs_ and wt_H2B-2FPs_ pachytene nuclei revealed a strong reduction in chromatin compaction levels in the absence of *set-2*, as indicated by a longer GFP-H2B fluorescence lifetime (Figure 1A), and a reduced mean-FRET efficiency (Figure 1B). From the measurement and spatial mapping of FRET in individual nuclei of wt_H2B-2FPs_ pachytene-stage cells, we observed discrete regions associated with distinct FRET efficiencies across individual nuclei (Figure 1A). As previously described (Llères *et al*, 2017), based on FRET quantification we arbitrarily defined several classes of FRET, from “sub-low FRET” to “high-FRET”, that correspond to different levels of nanoscale compaction. Interestingly, we observed that in the absence of *set-2*, the “intermediate-FRET” and “high-FRET” populations previously linked to heterochromatic states (Llères *et al*, 2017) were significantly reduced compared to wild type (Figure 1C), while the “sub-low-FRET” chromatin class associated with more accessible chromatin was increased. These results suggest that the absence of SET-2 results in changes in nanoscale chromatin structure in the germline, and more specifically disrupts highly compacted states.

### Loss of *set-2* enhances defects in germline chromatin organization resulting from condensin-II knock-down

Proper chromatin organization is required for chromosome segregation, and its disruption can result in chromosome segregation defects in mitosis and meiosis (Potapova & Gorbsky, 2017). These defects were not found in *set-2* mutant germlines or embryos at any of the temperatures tested (Beurton *et al*, 2019; Herbette *et al*, 2017). However, at the permissive temperature (20°C), endoreplicated intestinal cells of adult animals showed chromosome segregation defects very similar to those reported in condensin-II mutants (Csankovszki *et al*, 2009). This suggested that in *set-2* mutants, subtle defects in chromatin organization may arise that become apparent only in a sensitized background in which chromatin structure is further perturbed.

Condensins are major contributors to chromosome structure and organization (Hirano, 2012). Metazoans contain two types of condensin complexes (I and II) that share a heterodimer of two ‘structural maintenance of chromosomes’ (SMC) proteins, SMC2 and SMC4, and are distinguished by three unique CAP (chromosome-associated polypeptide) proteins named CAPD, CAPG and CAPH (Ono *et al*, 2004). Uniquely, *C. elegans* has an additional complex, condensin-IDC, which contributes exclusively to dosage compensation in somatic cells (Albritton & Ercan, 2018). KLE-2, HCP-6 and CAPG-2 are condensin-II specific subunits, CAPG-1, DPY-26 and DPY-28 are common to the two condensin-I complexes, whereas DPY-27 is specific to condensin-IDC (Figure 2A) (Csankovszki *et al*, 2009). In *C. elegans,* condensin-II associates with sister chromatids in meiosis and mediates their compaction and resolution (Chan *et al*, 2004; Mets & Meyer, 2009).

**Figure 2.**
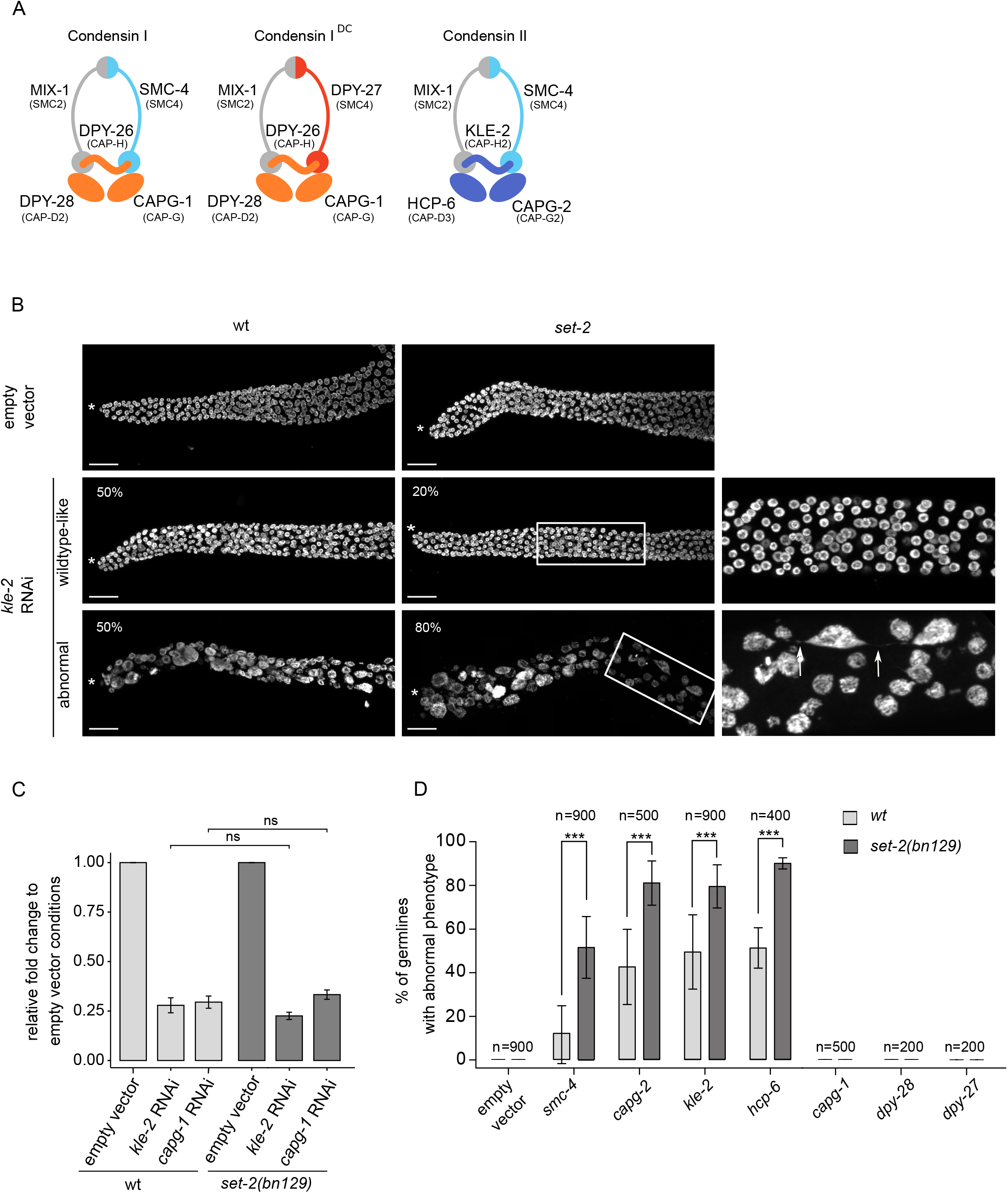
*set-2* inactivation enhances condensin-II depletion phenotypes. (**A**) *C. elegans* condensin subunits and their vertebrate homologs. (**B**) Confocal images of distal germline region from wildtype and *set-2(bn129)* animals treated with empty vector or *kle-2* RNAi. (*) marks distal end of the gonad. Representative images show examples of “wildtype-like” and “abnormal” phenotypes, with their presence indicated as percentage (%) of total (n=900, from 9 independent biological replicates) (scale bar, 20 μm). Arrow indicates the presence of chromatin bridge (scale bar, 10 μm). Images correspond to a Max intensity projection using using Fiji. (**C**) *kle-2* and *capg-1* mRNA levels in wildtype and *set-2(bn129)* mutant animals after RNAi directed against the respective genes. Relative fold change was calculated with respect to empty vector condition, following normalization with *pmp-3* and *cdc-42*. [*] p<0.05 t-test. (**D**) Percentage of germlines with “abnormal” phenotype after RNAi directed against condensin-II (*smc-4*, *capg-2*, *kle-2* and *hcp-6*), condensins I (*capg-1, dpy-28*) and condensin IDC (*dpy-27*) in wildtype or *set-2(bn129)* mutants. n= number of animals scored from 9 independent experiments for *smc-4* and *kle-2*, 4 for *hcp-6*, and 5 for *capg-1* and *capg-2*, and 2 for *dpy-28* and *dpy27*. All scoring was performed in blind. [***] p<0.001 (t-test).

Because depletion of condensin-II subunits results in sterility, we could not study the interaction of single condensin-II mutants with *set-2*. Instead, we knocked-down different subunits in wildtype and s*et-2* mutant animals by growing animals from the L1 larval stage to adulthood on condensin RNAi feeding plates, followed by scoring of DAPI stained germlines by fluorescence microscopy (Figure 2B and D). We initially focused on *kle-2* and *capg-1* RNAi to knock-down condensin-II and condensin-I complexes, respectively. RT-qPCR analysis showed that RNAi treatments resulted in a similar decrease in transcript levels in both wildtype and *set-2* mutant animals (Figure 2C), confirming that the efficiency of RNAi was the same in both genetic contexts.

Condensin-II RNAi resulted in reduced fertility in both wildtype and mutant animals. Because many of these animals also showed defects in vulval development that prevented egg-laying, we were unable to use the number of progeny laid as a read-out of fertility in wildtype animals compared to mutants. We instead used visual scoring of the germlines, placing animals in one of two broad classes: “wildtype-like” or “abnormal” (Figures 2B and D). Germlines in the wildtype-like class consist of nuclei undergoing all stages of meiotic progression as in wild type, although the total number of germ cells is reduced, consistent with the severe under-proliferation observed in condensin-II mutants (Csankovszki *et al*, 2009). The second class defined as “abnormal” consists of severely disorganized germlines containing fewer and larger nuclei, often showing more intense DAPI staining (Figure 2B and D). Using a lacO/lacI-GFP system composed of a stably integrated lacO array and a lacI::GFP fusion protein able to bind LacO repeats (Gonzalez-Serricchio & Sternberg, 2006), we observed multiple spots in enlarged nuclei, revealing that these were aneuploid nuclei (Figure S3). The abnormal germline morphology of these mutants made it difficult to clearly distinguish different region of the germline, and individual cells could not be unequivocally assigned to a specific meiotic stage. Nuclei were sometimes connected by thin chromatin bridges (Figure 2B, arrow), consistent with the known involvement of condensin-II in chromosome segregation in the germline and soma (Csankovszki *et al*, 2009; Stear & Roth, 2002).

RNAi of the condensin-II subunit *kle-2* in wildtype animals resulted in a comparable number of germlines falling in the wildtype-like and abnormal class (Figure 2B). *kle-2*(RNAi) in *set-2* mutant animals resulted in similar phenotypes, but there were significantly more germlines showing an abnormal phenotype, representing 80% of all germlines in blind scoring experiments. Similar results were observed following RNAi knock-down of the other condensin-II specific subunits, *hcp-6* and *capg-2*, and of *smc-4*, common to both condensin I and II complexes. In all conditions, phenotypes were consistently and significantly more severe in *set-2* mutant animals than in wildtype animals (Figure 2D).

Knockdown of the condensin-I specific subunits *capg-1* and *dpy-28*, or the condensin-IDC subunit *dpy-27* (Csankovszki *et al*, 2009; Chan *et al*, 2004; Mets & Meyer, 2009), did not produce any apparent germline phenotype, either alone, or in the *set-2* mutant background (Figure 2D). The effectiveness of *capg-1*, *dpy-28* and *dpy-27* RNAi was confirmed by scoring the associated dumpy (Dpy) phenotype in wildtype and mutant animals (Figure S4; www.wormbase.com).

In summary, germline phenotypes resulting from condensin-II knock-down are significantly and reproducibly more severe in the absence of *set-2*, consistent with cooperation between SET-2 and condensin in shaping meiotic chromosomes.

### Loss of *set-2* differentially affects germline and somatic phenotypes of the condensin-II mutant *hcp-6(mr17)*

To validate the above results based on RNAi knock-down, we used *mr17*, a hypomorphic allele of the condensin-II subunit *hcp-6* that carries a missense mutation resulting in temperature-sensitive embryonic lethality (Stear & Roth, 2002). In their previous analysis, the authors identified it as a G to A transition at nucleotide position 3073 within the coding region, converting glycine 1024 to glutamic acid (Stear & Roth, 2002). We could not confirm the presence of this mutation by resequencing. Instead, we identified a glycine to glutamic acid substitution at amino acid 683 within the HEAT repeat of HCP-6 in 3 independent experiments (Figure 3A). The reasons for this discrepancy are not clear, but may reflect errors in sequencing in previous experiments.

**Figure 3.**
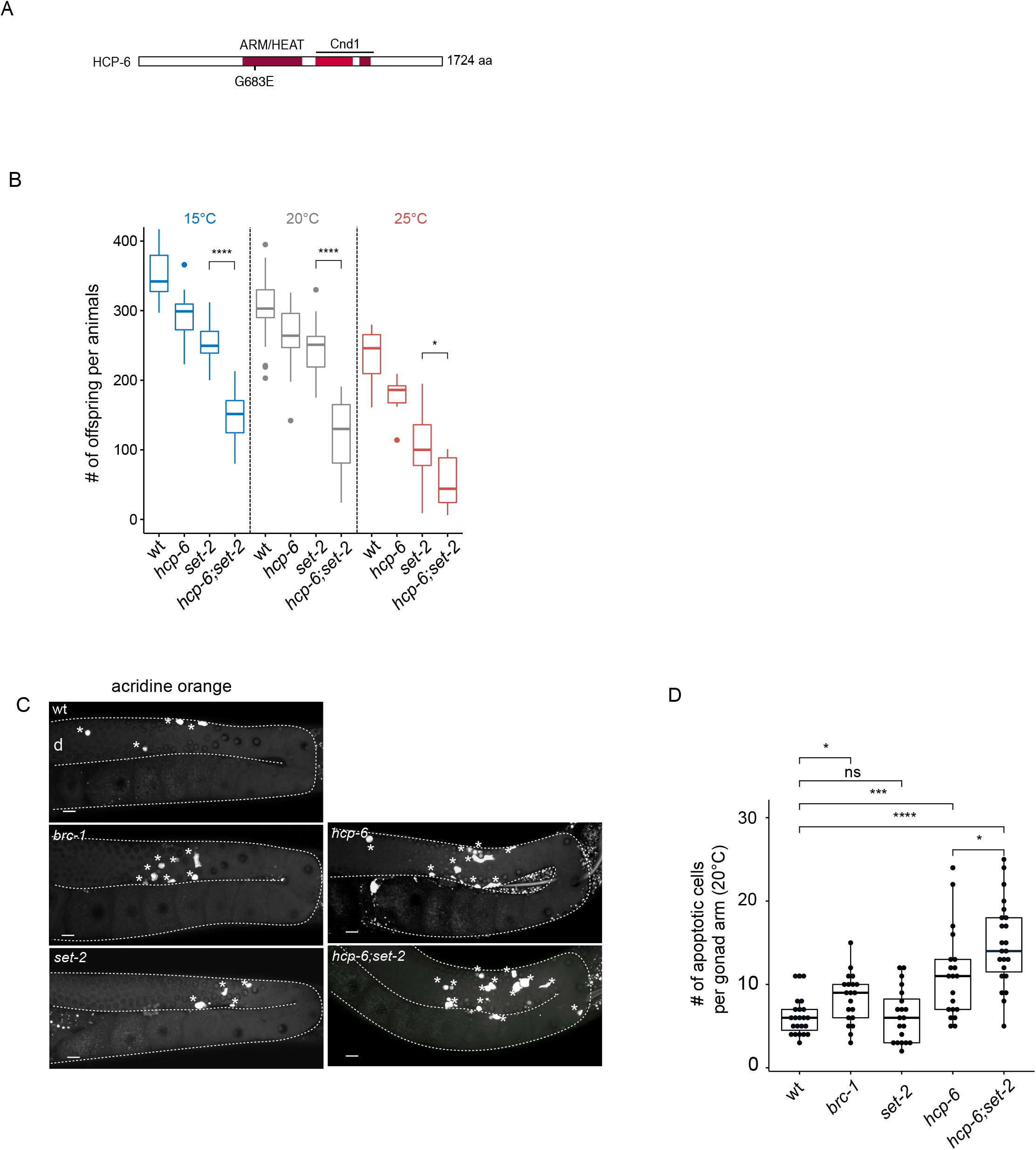
Enhancement of *hcp-6(mr17)* phenotypes in *set-2(bn129)* mutant animals. (**A**) Schematic diagram of HCP-6 protein and position of the *mr17* mutation. Conserved ARM/HEAT and Cnd1 (Condensin complex subunit 1) domains are highlighted in red. (**B**) Brood size per animals at indicated temperatures (15°C, 20°, 25°C), (n=11 animals per genotypes at 15°C and 25°C, and 33 at 20°C). [***] p < 0.001, [*] < 0.05 (t-test adjusted for multiple comparison with the Bonferroni method). (**C**) Representative confocal images of germlines stained by orange acridine. Asterisks indicate apoptotic cells, (d) indicates distal region; orientation is the same for all gonads. (**D**) quantification of the number of apoptotic cells in the germline of animals switched at 20°C for 24h (n>20 gonads per genotypes). A Wilcoxon test was performed after a significant difference with a Kruskal Wallis test, [ns] non significative difference, [*] p <0.05, [***] p <0.001, [****] p <0.0001.

Because the germlines of *hcp-6(mr17)* single and double mutants are severely disorganized at 25°C (Figure S5 A), we were unable to visually score distinct phenotypic classes in single compared to *hcp-6(mr17);set-2* double mutants, as we did for the RNAi experiments. However, we observed that *hcp-6(mr17)* mutants showed a significant reduction in the number of progeny laid at all temperatures tested (15°, 20°, 25°C) (Figure 3B), consistent with the essential role of condensin-II in germline fertility (Csankovszki *et al*, 2009). Using total brood size as a read-out of germline health, we found that *set-2* single mutants have reduced fertility, as previously described (Xiao *et al*, 2011). Brood size was further and significantly reduced in *hcp-6(mr17);set-2* double mutants compared to either of the single mutants at all temperatures (Figure 3B). Furthermore, in all cases, the *hcp-6(mr17);set-2* double mutant phenotype was most severe at the non-permissive temperature of 25°C. *hcp-6(mr17);set-2* double mutants laid a mean of 30 embryos per animal at 25°C, compared to 250-300 for wild type, showing that germline function is severely impacted in these animals.

Apoptosis eliminates defective germline nuclei in the *C*. *elegans* germline (Gumienny *et al*, 1999) and could contribute to the reduced brood size of *hcp-6* mutant animals. Using the dye acridine orange to mark apoptotic cells (Gumienny *et al*, 1999), we observed as expected an increase in apoptosis in *brc-1/*BRCA1 mutants (Boulton *et al*, 2004), while *set-2* single mutants were not affected (Figures 3C, D, (Herbette *et al*, 2017). The number of apoptotic corpses increased in *hcp-6* mutants, with a significant further increase in *set-2*;*hcp-6(mr17)* double mutants. The elimination of defective germline nuclei by apoptosis may therefore contribute to the reduced fertility of these animals (Figures 3C and D). Alternative, or in addition, a decrease in the mitotic stem cell population may contribute to reduced fertility. Altogether, our results are consistent with RNAi knock-down experiments and support functional cooperation between condensin-II and *set-2* in the germline.

The *hcp-6(mr17)* allele also allowed us to explore whether *set-2* genetically interacts with condensin-II in the soma. L4 animals raised at the permissive temperature of 15°C were shifted to 20°C and allowed to develop into adults, and their progeny scored. Under these conditions, the *hcp-6(mr17)* mutation resulted in greater than 85% embryonic lethality, which was reduced to 65% in *set-2*;*hcp-6(mr17)* double mutants (Figure S5 B). Absence of *set-2* did not reduce the embryonic lethality of *hcp-6(mr17)* mutants by suppressing cell division defects in these embryos, because *set-2*; *hcp-6(mr17)* surviving animals developed into adults showing phenotypes commonly associated with cell division defects, including uncoordinated behavior (unc) and sterility (O’Connell *et al*, 1998). Together, these results suggest that *set-2* and condensin-II subunit *hcp-6* have distinct functional relationships in the soma and germline.

### Functional links between SET-2 and chromosome structural protein TOP-2 in germline organization

Topoisomerase II is another major component of mitotic chromosome. When temperature-sensitive topoisomerase II mutants are used to bypass its essential requirement in mitosis, defects in chromosome condensation and segregation are also observed in meiosis (Marchetti *et al*, 2001;Li *et al*, 2013;Gómez *et al*, 2014;Hughes & Hawley, 2014). To explore whether *set-*2 also genetically interacts with *top-2* to ensure proper chromosome condensation in the germline, we constructed *set-2;top-2* double mutants using a recently described allele, *top-2 (it7)*, that results in a temperature-sensitive chromosome segregation defect in male spermatogenesis (Jaramillo-Lambert *et al*, 2016).

We observed that *top-2(it7)* hermaphrodites that developed from animals shifted to the non-permissive temperature (24°C) at the L1 larval stage also had defects in germline organization. Most adults contained normal germline arms whose size and developmental transitions were comparable to wild type (Figure 4A). The remaining animals presented either a smaller germline, such that germline bends were premature, or a germline atrophy phenotype with only a small population of mitotic germ cells. Notably, animals that displayed germline atrophy were mostly devoid of germ cells at various stages of meiosis (Kimble & Crittenden, 2005), and nuclei were larger in size. Short and atrophied germlines were significantly more abundant in *top-2 (it7);set-2* double than in *top-2* single mutants, accounting for 65% of all germlines scored (n=400) (Figure 4B). In addition, a minor fraction of germlines consisted of only mitotic germ cells (tumorous phenotype, data not shown). The extensive disorganization and decompaction of germline chromatin in *top-2 (it7);set-2* double mutants suggests that chromatin architecture is significantly altered in the germline of these animals.

**Figure 4.**
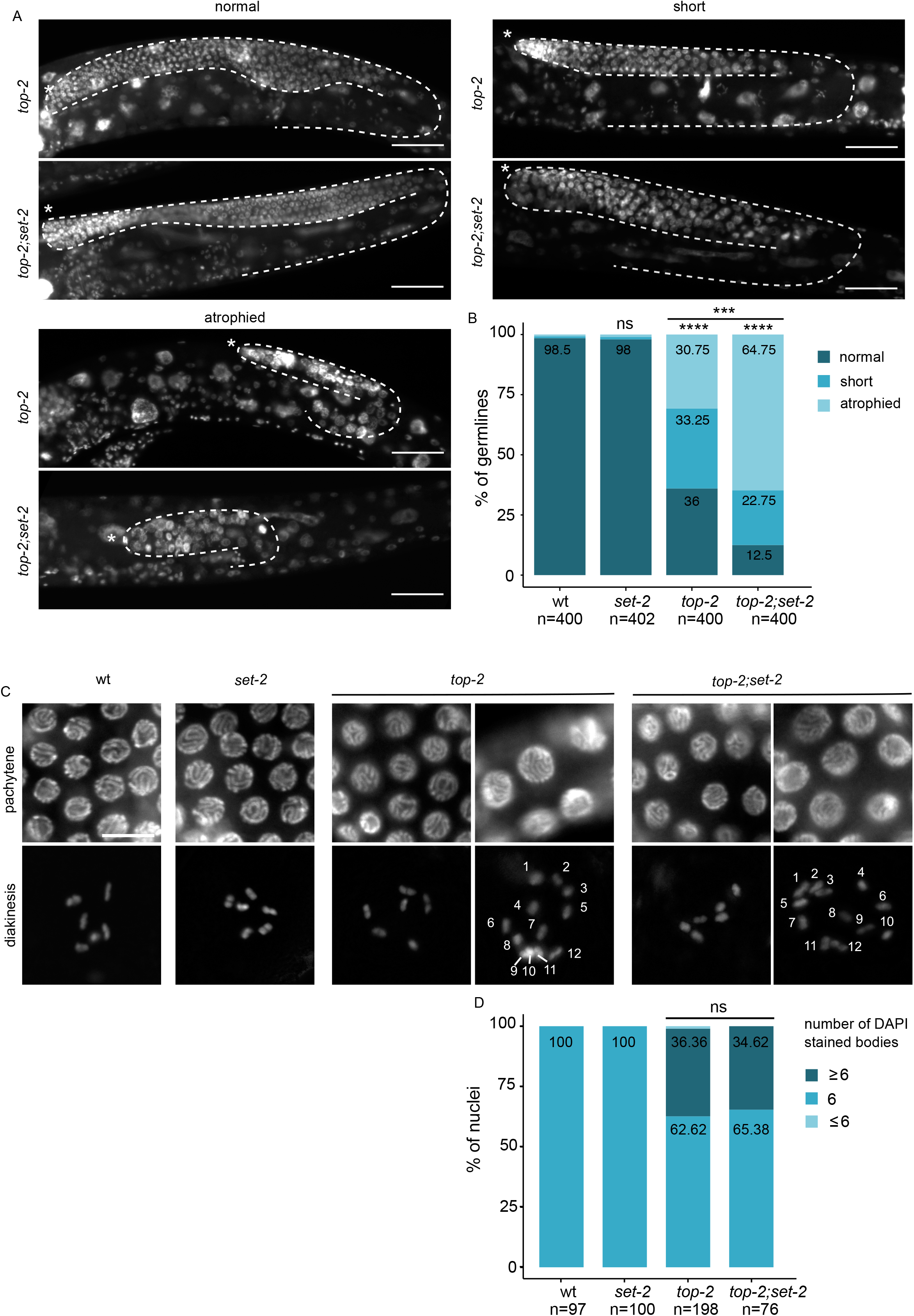
Enhancement of the *top-2* mutant phenotype in absence of *set-2*. (**A**) Representative images of DAPI stained adult germlines showing different phenotypic classes. (*) marks distal end of the gonad (**B**) Scoring of phenotypic classes. Animals were shifted to 24°C at the L1 stage and allowed to develop to adulthood. Germlines categories are as defined in Materials and Methods. **** p<0.0001, *** p<0.001 significant difference between mutant backgrounds using chi-square test and FDR correction (ns: non-significant). Scale bar, 50 μm. (**C**) Enlargement of pachytene and diakinetic nuclei from wildtype or mutant animals. For *top-2* single and *top-2;set-2* double mutants, representative nuclei from normal (left) and short (right) germlines are shown. Nuclei containing more than 6 DAPI stained bodies were observed in both “short” and “normal” germlines. (**D**) Scoring of aneuploid nuclei from cells in diakinesis (Scale bar, 10 μm).

In wildtype oocytes, individual chromosomes appear as 6 condensed DAPI stained structures (Kimble & Crittenden, 2005). Consistent with the established role of topoisomerase II in sister chromatid segregation (Liang *et al*, 2015; Nagasaka *et al*, 2016; Oliveira *et al*, 2010; DiNardo *et al*, 1984), *top-2(it7)* mutants had a significant number of oocytes (67%) with more than 6 DAPI-stained bodies, and a smaller number (33%) with fewer (Figure 4C and D). A similar phenotype was observed in *top-2(it7);set-2* double mutants, suggesting that topoisomerase II acts independently of SET-2 in sister chromatid cohesion. Together, these results are consistent with a role for SET-2 in chromatin organization of pachytene chromosomes together with condensin and Topo II, two major components of chromosome architecture.

### *set-2* inactivation does not affect germline expression of condensins, topoisomerase, or somatic genes

Previous transcriptome profiling of *set-2* mutant germlines by microarray analysis revealed upregulation of many somatic genes (Robert *et al*, 2014). However, these experiments were carried out on mutant animals maintained in the homozygous state and likely to have incurred additional changes in gene expression not directly related to the absence of SET-2. We therefore repeated transcription profiling by RNA-sequencing (RNA-seq) of dissected gonads from *set-2/set-2* homozygotes derived from *set-2/+* heterozygous mothers. Setting the p value at ≤0,05, 1670 genes were found to be down-regulated, and 1553 were up-regulated (Table S1). Genes expressed in the germline (pregametes and oocytes) were overrepresented in the set of upregulated genes (Figure S6). In contrast to our previous experiments, somatic genes were not over-represented in our data set, suggesting that their misregulation is a secondary consequence of propagating *set-2* mutant homozygotes several generations at 20°C (Figure S6). We also did not observe mis-regulation of either condensins or topoisomerase. Altogether, our analyses suggests that the meiotic chromosome organization defects we observe in *set-2* mutant germlines in combination with condensin and topoisomerase depletion are unlikely to simply reflect decreased expression of these genes, or a change in cell fate resulting from the expression of somatic genes. However, because transcription has emerged as a major contributor to chromatin architecture in many eukaryotes (Rowley *et al.*, 2017; van Steensel & Furlong, 2019), altered gene expression patterns in *set-2* mutant germlines may contribute to changes in chromatin architecture in these mutants.

### Loss of COMPASS targeting subunit CFP-1 results in similar chromatin organization defects as SET-2 inactivation

To establish whether SET-2 contributes to germline chromatin organization in the context of COMPASS, we next asked whether other subunits of the complex also enhance the germline defects resulting from condensin-II knockdown. CFP-1 is the chromatin-targeting subunit of COMPASS (Clouaire *et al*, 2012). As observed for *set-2*, inactivation of *cfp-1* using either the deletion allele *tm6369* or RNAi resulted in a strong decrease in H3K4me3 in both the germline and soma (Chen *et al*, 2014; Li & Kelly, 2011; Simonet *et al*, 2007; Xiao *et al*, 2011) Figure S7 A). The number of animals with an abnormal germline phenotype following RNAi of *smc-4*, targeting condensin-I and -II, or *kle-2*, targeting condensin-II only, was largely increased in *cfp-1* mutants compared to wild type (Figure 5A). FLIM-FRET analysis of *cfp-1* mutants expressing H2B-2FP showed a drastic nanoscale decompaction of pachytene chromatin (Figure 5B). This further supports a role for COMPASS in chromosome organization.

**Figure 5.**
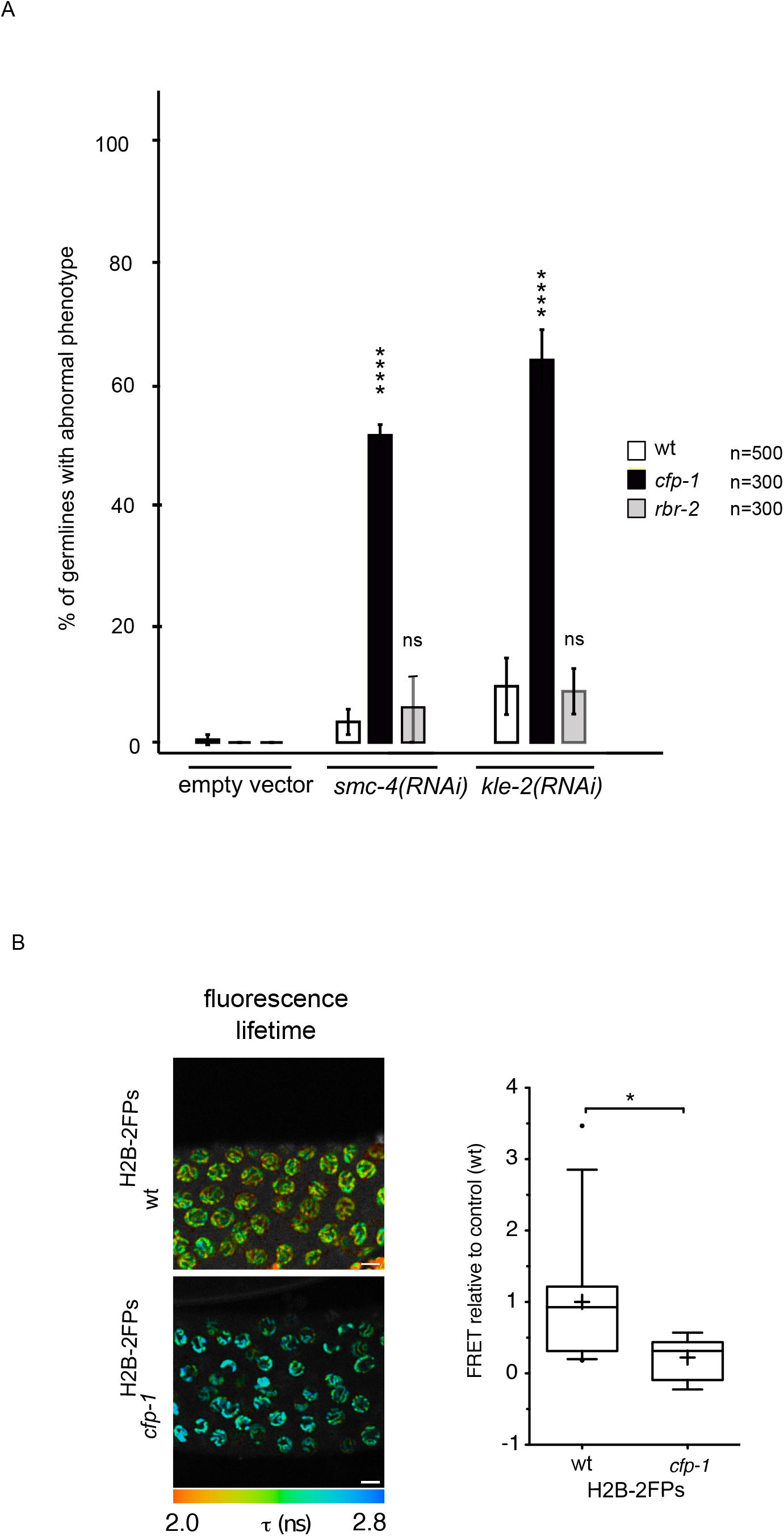
Inactivation of COMPASS targeting component *cfp-1* mimics *set-2* inactivation. (**A**) Percentage of germlines with “strong” phenotype after RNAi directed against condensin-II (*smc-4* or *kle-2*) in wildtype, *cfp-1(tm6369)* and *rbr-2(tm1231)* mutants. n= number of animals scored from at least 3 independent experiments. p <0.05, [***] p <0.001, [****] p <0.0001 (t-test); ns, not significant. (**B**) Decreased chromatin compaction in *cfp-1* mutant germlines. Spatial distribution of the mean fluorescence lifetime is shown for wildtype (wt _H2B-2FPs_) or *cfp-1* _H2B-2FPs_ mutant animals. Fluorescence lifetime values (τ) ranging from 2.0 ns to 2.8 ns are represented using a continuous pseudo-color scale. Scale bars, 10 μm. Statistical analysis of the FRET efficiency relative to control wt is presented as box- and-whisker plots. Mean FRET value is indicated by a cross in each box. *p < 0.05 (two-tailed unpaired t test). n=13 gonads for wt, n=11 gonads for *cfp-1*.

H3K4me3 is removed by the well-conserved lysine demethylase RBR-2/KDM5 (Christensen *et al*, 2007; Alvares *et al*, 2014). Like *set-2*, *rbr-2* is required to maintain germline immortality at high temperatures (Alvares *et al*, 2014). We found that absence of RBR-2 activity in the *rbr-2(tm1231)* deletion allele (Alvares *et al*, 2014) did not result in the enhancement of germline defects resulting from either *kle-2* or *smc-4* (RNAi) (Figure 5A). qRT-PCR analysis confirmed that although the overall efficacy of RNAi varied between independent experiments, RNAi efficacy was comparable in wildtype, *cfp-1* and *rbr-2* mutants within the same experiment (Figure S7 B). Furthermore, the percentage of animals with a strong phenotype was similar in all three experiments (Figure S7 C), consistent with depletion of condensin-II below a threshold level being sufficient to provoke defects in chromosome organization (Samejima *et al*, 2012). Therefore, contrary to COMPASS inactivation, increasing H3K4 methylation levels in *rbr-2* mutants has no obvious impact on germline chromatin organization.

## Discussion

Using three different experimental approaches, we found that in the *C. elegans* germline the COMPASS H3K4 methyltransferase complex contributes to the organization of chromosome architecture. First, using quantitative tagged-histone FLIM-FRET analysis we discovered that chromatin compaction at the nucleosomal level is reduced in live animals that lack the COMPASS subunits SET-2 or CFP-1. Second, we demonstrate that defects in the organization and compaction of germline nuclei following knockdown of condensin-II subunits or topoisomerase II are strongly enhanced in the absence of COMPASS. Third, using the number of progeny as a measure of germline health, we demonstrate that *set-2* inactivation exacerbates the fertility defects associated with a hypomorphic allele of the condensin-II subunit *hcp-6*. Combined, these findings evoke a role for COMPASS in shaping chromosome structure in *C. elegans.*

Our transcriptomic analysis of germ cells lacking *set-2* did not reveal significant downregulation of condensins, topoisomerases, or other genes encoding chromatin factors with known function in *C. elegans* chromatin organization, making it unlikely that COMPASS alters chromosome structure through misregulation of any of these genes. We consider a direct effect of COMPASS on condensin-II also unlikely, since we found no evidence of a physical interaction between the two in extensive proteomics analysis (Beurton *et al*, 2019) and data not shown). Rather, based on our data we favor a more direct role for COMPASS in modifying meiotic chromosome organization.

A current model proposes that condensin complexes topologically shape mitotic chromosomes through a loop extrusion process (Ganji *et al*, 2018), with condensin-I and -II forming arrays of helical consecutive loops in mitotic cells (Gibcus *et al*, 2018). However, the observation that chromosomes still maintain a certain degree of structure in the absence of both condensin-I and -II suggests that additional mechanisms and factors may be involved, including histone modifying complexes and the associated modifications (Samejima *et al*, 2012; Wilkins *et al*, 2014; Kruitwagen *et al*, 2015; Markaki *et al*, 2009; Zhiteneva *et al*, 2017;Vagnarelli *et al*, 2006). Using the same FLIM-FRET imaging approach implemented here, we previously showed that condensin complexes contribute to the nanoscale compaction of chromatin in the *C. elegans* germline (Llères *et al*, 2017). Depletion of condensin-I affected both highly- and lowly-compacted regions, whereas depletion of condensin-II only affected highly compacted regions (Llères *et al*, 2017). By using FRET imaging to quantify proximity between fluorescently labelled nucleosomes, in the current study we observed that *set-2* controls the same compacted chromatin states as condensin-I (Llères *et al*, 2017). Because FRET in our system most likely results from clustering of distant regions from the same or different chromosomes, our findings most likely imply that *set-2*, and to a larger extent COMPASS, contributes to the formation of chromatin loops to establish the proper structural organization of meiotic chromosomes.

In mammals, partial inactivation of condensin-II results in relatively moderate defects in chromosome structure, in contrast to the more severe defects resulting from its complete inactivation (Ono *et al*, 2004; Vagnarelli *et al*, 2006; Hudson *et al*, 2003). By employing a similar approach, we found that condensin-II RNAi results in less compacted pachytene chromosomes. Absence of COMPASS subunits *set-2* or *cfp-1* significantly aggravated this defect, suggesting that SET-2 cooperates with condensin-II for proper meiotic chromatin architecture. We further confirmed that COMPASS and condensin-II functionally interact by showing that reduced fertility of the hypomorphic allele *hcp-6*(*mr17*) is aggravated in the absence of *set-2*. Interestingly, by resequencing the *hcp-6*(*mr17*) allele, we identified the mutation as a substitution of glycine with glutamic acid within one of the α-helical HEAT repeats of HCP-6. Although the role of the HEAT-repeat subunits, which are unique to eukaryotic condensins, remains largely unknown (Yoshimura & Hirano, 2016), our results suggest that they are of functional importance.

Interestingly, we also found that *set-2* partially suppressed the embryonic lethality of *hcp-6*(*mr17*) mutants at the non-permissive temperature. This effect may be due to elimination of defective germ cells by increased apoptosis in *set-2;hcp-6* double mutant germlines, thereby improving the quality of surviving oocytes with respect to the *hcp-6* single mutant. We note that mutations in the BRCA1 homologs *brc-1* or *brd-1* are also able to partially suppress the embryonic lethality of *hcp-6*(RNAi) animals, possibly reflecting a role for BRC-1/BRD-1 in the formation of toxic chromatin bridges when chromosome condensation is defective (Hong *et al*, 2016). Our results suggest that SET-2 may also be involved in a similar process in embryos.

Further insight into a likely role of SET-2 in higher-order chromatin organization comes from the observation that *set-2* inactivation also enhanced the chromatin organization defects of *top-2* conditional mutants. Topoisomerase-II contributes to proper mitotic condensation and structure in all species studied, including *C. elegans* (Cuvier & Hirano, 2003; Petrova *et al*, 2013; Sakaguchi & Kikuchi, 2004; Samejima *et al*, 2012; Shintomi *et al*, 2015; Uemura *et al*, 1987). In *C. elegans*, TOP-2 localization along mitotic chromosomes is thought to constrain chromosome length by modulating chromatin loops (Ladouceur *et al*, 2017). Topoisomerase II also localizes along the chromosome axes of meiosis I chromosomes, both in yeast and mammals (Gómez *et al*, 2014; Klein, 1992; Li *et al*, 2013), and it may play a similar structural role in organizing *C. elegans* germline nuclei in cooperation with SET-2.

The enhancement of the condensin-II and topoisomerase knock-down phenotypes by COMPASS inactivation suggests at least partially independent, cumulative, effects on chromosome organization (Figure 6). Meiotic chromosomes from yeast to mammals are organized as linear loop arrays around a proteinaceous chromosome axis (Blat *et al*, 2002; McNicoll *et al*, 2013; Panizza *et al*, 2011; West *et al*, 2019), and recent super-resolution microscopy studies on mouse oocytes show that H3K4me3 emanates radially in similar structures that contribute to the shaping of meiotic chromosomes (Prakash *et al*, 2015). In mouse spermatocytes, strong clustering of highly transcribed loci is observed and is thought to reflect local interactions between linear loops, as well as between loci on homologs (Patel *et al*, 2019). Based on these observations, we suggest that the presence of COMPASS at transcription sites could contribute to the organization of chromatin in these clusters. This could take place through the recruitment of an H3K4me3 reader, as described for the NCAPD3 subunit of condensin-II (Yuen *et al*, 2017), through modifications in the chromatin landscape brought about by *set-2* dependent changes in gene expression (Rowley *et al*, 2017; van Steensel & Furlong, 2019), or through the recruitment of additional proteins (Beurton *et al*, 2019). Finally, given the recent implication of the nuclear RNAi pathway and the MORC-1 gene silencing protein in germline chromatin organization (Kim *et al*, 2019; Weiser *et al*, 2017), additional interactions between COMPASS and these pathways may be involved.

**Figure 6.**
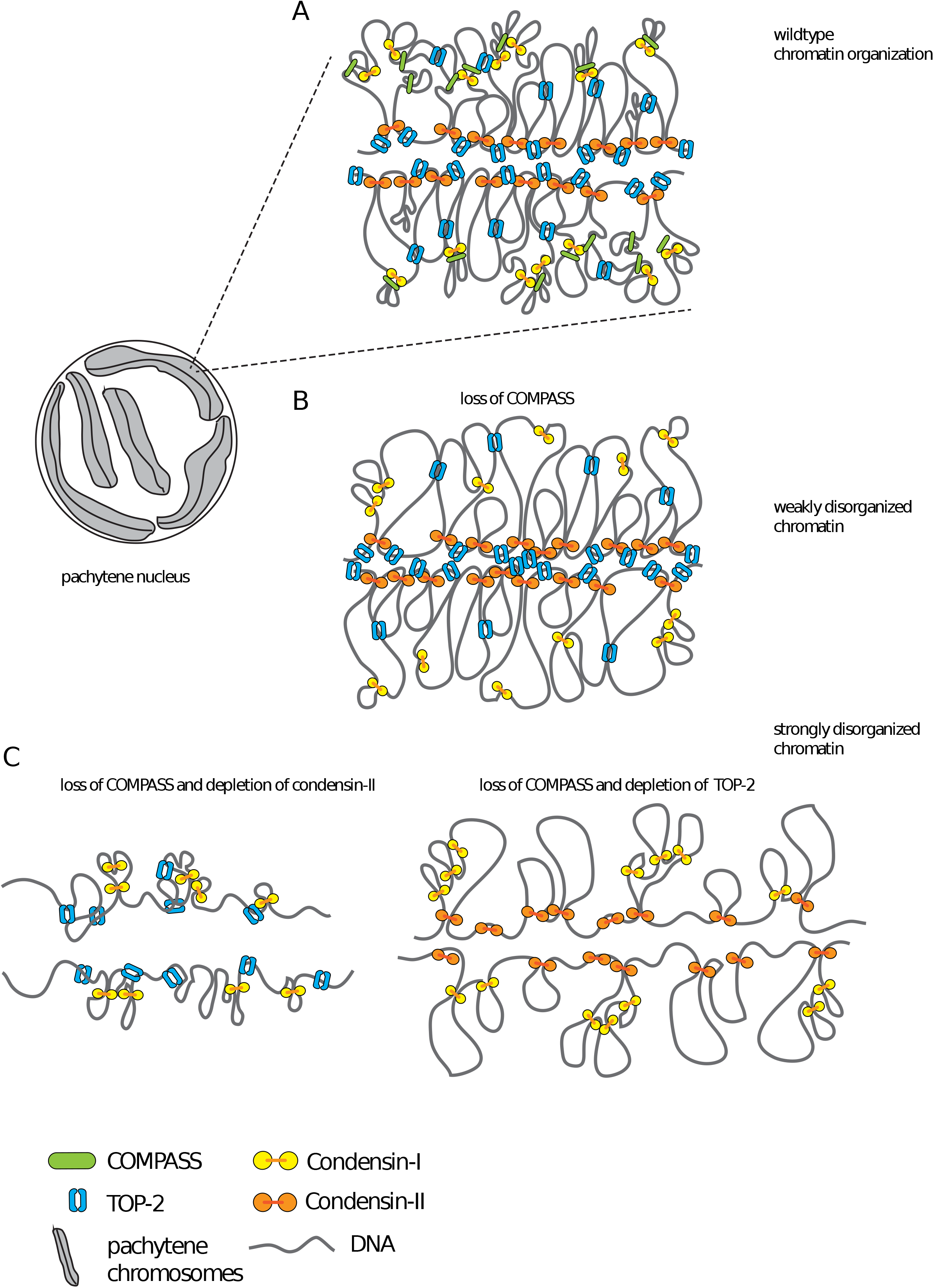
Working model for cooperation between COMPASS, condensin-II and Topoisomerase-II in chromosome organization. In wild type, proper chromosome compaction results from the activity of condensin-I (connected yellow circles), condensin-II (connected orange circles) and Topoisomerase-II (blue broken ring). The concerted action of condensins results in the formation of arrays of helical loops, with condensin-II generating outer loops and condensin-I forming inner loops (Gibcus *et al*, 2018). Topoisomerase-II may contribute to compaction by modulating chromatin loops (Ladouceur *et al*, 2017), or actively introducing self-entanglement in DNA (Piskadlo & Oliveira, 2016). COMPASS may mediate interactions between loops, possibly by contributing to the clustering of transcribed loci (Patel *et al*, 2019). The absence of COMPASS results in subtle defects in chromosome organization (weakly disorganized chromatin), but overall chromosome architecture is maintained by the action of condensins, Topoisomerase-II and additional proteins.Partial depletion of condensin-II or Topoisomerase-II in the absence of COMPASS results in cumulative defects in chromosome organization.

Our studies highlight an important role for the evolutionarily conserved COMPASS complex in chromosome organization in the *C. elegans* germline. Deciphering the mechanism whereby chromatin-associated factors and histone post-translational modifications affect global regulation of chromatin architecture in meiosis will be an important area of future study. Given the remarkable morphological similarities between chromosomes in meiotic prophase and early mitotic prophase (Alavattam *et al*, 2019; Jabbari *et al*, 2019; Schalbetter *et al*, 2019; Wang *et al*, 2019b; MacGregor *et al*, 2019; Gibcus *et al*, 2018), our germline findings may have implications for our understanding of mitotic chromosome condensation as well.

## Methods

### Nematode maintenance and strains

Unless otherwise noted, animals were propagated under standard conditions at 20°C or 15°C (Brenner, 1974) on NGM plates (Nematode Growth Medium) seeded with the *Escherichia coli* strains OP50 or HT115 for RNAi experiments. N2 Bristol was used as the wildtype control strain. Strains used were as follows:

*hcp-6(mr17)* I (PFR656), *set-2(bn129)* III/qC1 (PFR510), *hcp-6(mr17)* I; *set-2(bn129)* III (PFR651), *cfp-1(tm6369)* IV (PFR588), *brc-1(tm1145)* III (DW102), *rbr-2(tm1231)* IV (PFR394), oxIs279[Ppie 1::GFP::H2B + *unc-119*(+)] II; *unc-119(ed3)* III (EG4601), oxI*s279*[P*pie-1*::GFP::H2B + *unc-119*(+)] II; *set-2(bn129)* III (PFR326), *oxIs279*[P*pie-1*::GFP::H2B + *unc-119*(+)] II; *cfp-1(tm6369)* IV (PFR667), oxSi487 [mex-5p::mCherry::H2B::tbb-2 3’UTR::*gpd-2* operon::GFP::H2B::*cye-1* 3’UTR + *unc-119*(+)] II; *unc-119(ed3)* III (EG6787), *oxSi487* [mex-5p::mCherry::H2B::tbb-2 3’UTR::*gpd-2* operon::GFP::H2B::cye-1 3’UTR + *unc-119*(+)] II; *set-2(bn129)* III (PFR659), oxSi487 [*mex-5*p::mCherry::H2B::*tbb-2* 3’UTR::*gpd-2* operon::GFP::H2B::*cye-1* 3’UTR + *unc-119*(+)] II; *cfp-1(tm6369)* IV (PFR666), *unc-119*(ed3) III; *top-2(it7)* II (PFR704), *top-2(it7)* II; *set-2(bn129)* III (PFR705)

### Worm Preparation for live-imaging

For FRAP and FLIM-FRET acquisitions, single worms (24 hours post-L4 stage) from an unsynchronized population were picked to an unseeded 1xNGM plate to wash off bacteria and were subsequently transferred onto a glass slide in a drop of egg buffer (118 mM NaCl, 48 mM KCl, 2 mM CaCl_2_*2H_2_O, 2 mM MgCl_2_*6H_2_O, 25 mM HEPES pH 7.3). Worm gonads were extruded by microdissection using a 23G syringe and immediately covered with a coverslip, sealed with nail varnish.

### FLIM-FRET data acquisition

FLIM-FRET measurements were carried out on wt, *set-2(bn129) and cfp-1(tm6369)* mutant strains GFP-H2B (donor alone: GFP-H2B protein) and H2B-2FPs (donor and acceptor: GFP-H2B and mCherry-H2B). FLIM was performed using an inverted laser scanning multi-photon LSM780 microscope (Zeiss) equipped with an environmental black-walled chamber. Measurements were performed at 20°C with a 40x oil immersion lens, NA 1.3 Plan-Apochromat objective, from Zeiss. Two-photon excitation was achieved using a tunable Chameleon Ultra II (680–1,080 nm) laser (Coherent) to pump a mode-locked, frequency-doubled Ti:sapphire laser that provided sub-150-fs pulses at an 80-MHz repetition rate. GFP and mCherry fluorophores were used as a FRET pair. The optimal two-photon excitation wavelength to excite the donor GFP was determined to be 890 nm (Llères *et al*, 2009). Laser power was adjusted to give a mean photon count rate of about 7.10^4^–10^5^ photons per second. Fluorescence lifetime measurements were acquired over 60 s. Detection of the emitted photons was achieved through the use of an HPM-100 module (Hamamatsu R10467-40 GaAsP hybrid photomultiplier tube [PMT]). and fluorescence lifetimes were calculated for all pixels in the field of view (256 x 256 pixels). The fluorescence lifetime imaging capability was provided by time-correlated single-photon counting (TCSPC) electronics (SPC-830; Becker & Hickl). TCSPC measures the time elapsed between laser pulses and the fluorescence photons. Specific regions of interest (e.g., full gonad or pachytene nuclei) were selected using SPCImage software (Becker & Hickl).

### FLIM-FRET Analysis

FLIM measurements were analyzed as described previously (Llères *et al*, 2017) using SPCImage software (Becker & Hickl). Briefly, FRET results from direct interactions between donor and acceptor molecules (T. & Förster, 1949) and causes a decrease in the fluorescence lifetime of the donor molecules (GFP). The FRET efficiency (*i.e*., coupling efficiency) was calculated by comparing the fluorescence lifetime values from FLIM measurements obtained for GFP donor fluorophores in the presence and absence of mCherry acceptor fluorophores. The FRET percentage images were calculated such as, *E FRET* = *1*-(τ*DA*/τ*D*), where τDA is the mean fluorescence lifetime of the donor (GFP-H2B) in the presence of the acceptor (mCherry-H2B) expressed in *C. elegans*_H2B-2FPs_, and τD is the mean fluorescence lifetime of the donor (GFP-H2B) expressed in *C. elegans*_GFP-H2B_ in the absence of the acceptor. In the non-FRET conditions, the mean fluorescence lifetime value of the donor was calculated from a mean of the τD by applying a mono-exponential decay model to fit the fluorescence lifetime decays.

### Condensin RNAi knockdown

Bacterial clones expressing RNA targeting condensin-I and -II subunit were from the *C. elegans* RNAi collection (Ahringer laboratory-Gene Service Inc). Inserts from each RNAi clone were amplified by PCR on isolated colonies, with a single primer in the duplicated T7 promoter (5’ TAATACGACTCACTATAGGG 3’), then sequenced using the primer 5’ GGTCGACGGTATCGATAAGC 3’. RNAi clones were cultured in LB liquid medium supplemented with 50μg/ml Ampicillin for 18h at 37°C, IPTG was then added (1mM final), and cultures grown an additional 2h30 at 37°C. NGM plates complemented with IPTG (1mM) were seeded with 300 μl of bacterial culture. Synchronized L1 were placed on RNAi plates and grown to adulthood.

### RNA isolation and qRT-PCR analysis

Synchronized L1 wildtype or *set-2(bn129)* mutant worms were grown on empty vector L4440 or *kle-2* or *capg-1* RNAi to adult staged worms at 20°C and harvested. Total RNA was isolated using NucleoZol (Macherey Nagel, #740404-200) and NucleoSpin (Macherey Nagel, #40609). RNA was reverse transcribed using cDNA transcriptor (Roche, #5893151001). Quantitative PCR analysis was performed on CFX Connect (Bio-rad CFX Connect) with SYBR Green RT-PCR (Roche, #4913914001). Melting curve analysis was performed for each primer set to ensure the specificity of the amplified product and with an efficiency of 2. *pmp-3* and *cdc-42* were used as the internal controls so that the RNA level of each gene of interest was normalized to the levels of *pmp-3* and *cdc-42*. qRT-PCR were performed on three biological replicas in technical duplicates. Statistical analysis was performed using an unpaired t-test. Primers used were:

*pmp-3*: 5’ GTTCCCGTGTTCATCACTCAT 3’ — 5’ ACACCGTCGAGAAGCTGTAGA 3’

*cdc-42*: 5’ CTGCTGGACAGGAAGATTACG 3’ — 5’ CTCGGACATTCTCGAATGAAG 3’

*kle-2:* 5’ GAGAAAACGGACAGCTCGTGTG 3’ — 5’ CGTCATATTCAGCTCCGAGGGT 3’

*capg-1:* 5’ TCGAATTGGCCAGTAGATGC 3’ — 5’ ACTGCAACAAGTCGGCATTC 3’

### Hoechst staining on dissected germlines

Germlines from condensin RNAi knock-down animals were dissected on L-polylysine coated slides in a drop of dissection buffer (0.4X M9 and Levamisole 20mM). After removing dissection buffer using a drawn capillary, gonads were fixed in 11μl of 3% paraformaldehyde for 5min. Slides were washed in 1X PBS 0.2% Tween 20 plus 5μg/ml Hoechst 33342 (Sigma Aldrich, #861405) for 10 min, then twice in 1X PBS 0.2% Tween 20 for 10 min, and mounted in mounting media (1X PBS, 4% n-Propyl-Gallate, 90% DE Glycerol). Z-stack images (0.25 μm slices) of germlines were acquired using a Zeiss LSM710 inverted confocal microscope with a 40X oil immersion objective.

### DAPI staining on whole animals

For scoring topo II mutant germlines, adult animals were stained as previously described (Kadyk & Kimble, 1998) with minor modifications. Briefly, animals were collected and washed once in 1X M9, fixed 15 min in −20°C methanol, and washed twice in 1X PBS with 0.1% Tween 20® (Sigma Aldrich, #P1379). 25μl Fluoroshield plus DAPI (Sigma Aldrich, #F6057) was added directly to 50μl of worm pellet, followed by mounting for fluorescent microscopy. Observation were made on an AxioImager A2 (Zeiss) with Plan Apochromat 63X/1.4 oil DIC or EC plan Neofluar 20X/0.5 objectives.

### Scoring of germline phenotypes

Blind scoring was carried out using AxioImager A2 (Zeiss) with EC plan Neofluar 20X/0.5 objectives. For condensin RNAi knock-down, the “abnormal phenotype” was defined as germlines containing fewer, abnormally sized and unevenly distributed nuclei, as well as macro nuclei with strong DAPI signal, as previously described (Csankovszki *et al*, 2009). “wildtype-like” includes germlines that mostly resembled wildtype and sometimes contained a few macro nuclei with strong DAPI signal. For each experiment, at least 200 germlines were scored for each genotype, and at least 3 independent biological replicates were performed. For experiments with the *top-2(it7)* allele, wildtype, *set-2(bn129)* and *top-2(it7)* single, and *set-2(bn129);top-2(it7)* double mutants were synchronized at the L1 stage at 15°C, then transfer on plates seeded with OP50 at 24°C and allowed to develop to adulthood. Adults were recovered in 1X M9 and DAPI stained as described in DAPI staining on whole animals. Germlines were place in phenotypic categories based on (Heestand *et al*, 2018). Data were collected from 3 independent experiments.

### Sequencing and mapping of the *hcp-6(mr17)* mutation

Genomic DNA from wildtype animals (N2) and from strains bearing the *hcp-6(mr17)* mutation was amplified using a high-fidelity polymerase (Phusion®, NEB #M0530S) and the following primer pairs:

5’ ATAGTCAACCTCGATTGCTGGCTG 3’ — 5’ GAGGGCGAATAAGTCTTCCGTAAG 3’

5’ GGAGTTTCTGCTGCCAGTAGTTAT 3’ — 5’ TGTGGATAAACGTGGCGATA 3’

5’ GATCGTTGGAGCGATTTACGGATC 3’ — 5’ TGTGGATAAACGTGGCGATA 3’

5’ GATCGTTGGAGCGATTTACGGATC 3’ — 5’ CTTTCTGGCATGTTCAGTGACGTC 3’

5’ GAAATCCCGAAGCAAGAGAG 3’ — 5’ GTCCATGTGAGATCCGATGAGT 3’

5’ GAAATCCCGAAGCAAGAGAG 3’ — 5’ CTTTCTGGCATGTTCAGTGACGTC 3’

5’ TGGCTTCACACCTTGATCTCGATG 3’ — 5’ TCTTCATCGTGACCAACTCCAACC 3’

5’ TCTCAACGTGGCATCTGAAG 3’ — 5’ GCGTGTCGACGAACAATAAC 3’

5’ GTTCGGAATGACGCAAAACT 3’ — 5’ CACAGTTTTCTCCGCATCAACATG 3’

5’ CACTGAAATGCGCCTTAATCCTCC 3’ — 5’ TGATATGGGAGGAGCTGTGAAGGA 3’

For each DNA fragment amplified by PCR, both forward and reverse primers were used in the sequencing reactions. The presence of the *mr17* mutation was confirmed in 7 independent sequencing reactions from independent isolates. 2 independent reactions from wildtype animals were used as reference.

### Brood size and embryonic lethality assays

To score fertility and embryonic lethality, 10 to 11 individual L4 hermaphrodites grown at 15°C were picked and transferred to individual plates at either 15°C, 20°C or 25°C. Animals were transferred on new plates until they stopped laying eggs, and the number of eggs on individual plates scored each day. After 24h, unhatched eggs and live progeny were scored. Embryonic lethality represents the number of unhatched eggs, divided by the total number of total eggs laid. Experiments were repeated 3 times each.

### Visualization of apoptotic cells in the germline

Acridine Orange (Sigma Aldrich #A9231) was used to visualized apoptotic cells in the germline of live animals as previously described (Papaluca & Ramotar, 2016). Briefly, L4 hermaphrodites grown at permissive temperature (15°C), were placed on NGM plates at the restrictive temperature of 20°C during 18h. 1ml of Acridine Orange diluted at a final concentration of 50μg/ml in M9 buffer was added to the plates and incubated for 2h in the dark. Stained animals were transferred to a fresh NGM plates seeded with OP50 and incubated for 2h in the dark in order to remove stained bacteria in the intestine. Animals were placed on 4% agar pad in a drop of 10mM levamisole (Sigma Aldrich #L9756) diluted in M9 buffer, a coverslip was placed on top and sealed with nailed polish. Z-stack images of the posterior gonad were acquired using a Zeiss LSM710 inverted confocal microscope with 40X oil Immersion objective. Z-stack of germlines were acquire every 0.5μm, images correspond to a projection using Max intensity method using Fiji (Schindelin *et al*, 2012). At least 20 gonads were imaged for each genotype.

## Declarations

All data in this manuscript is publicly available.

The authors declare that they have no competing interests.

## Authors’ contributions

FP and MH designed genetic experiments; MH performed and analyzed genetic assays; DL carried out FRAP and FLIM-FRET experiments; AB made genetic constructs and helped with FLIM-FRET acquisition and experimental design; VR helped with experimental design and helped carry out qRT-PCR analysis; MH and LG carried out immunofluorescence analysis; FP and DL wrote the paper; MH, FP and DL prepared figures, and all authors helped with editing; all authors read and approved the final manuscript.

## Acknowledgements

We thank Montpellier “Ressources Imageries” (MRI), CGC, which is funded by NIH Office of Research Infrastructure Programs (P40 OD010440) for strains, and the Mitani lab for deletion strains.

## Supplementary data

### H3K4me3 Immunostaining

Immunostaining was as previously described (Robert *et al*, 2014). Z-stack images of gonads were acquired using a Zeiss LSM710 inverted confocal microscope with 40X oil Immersion objective. Z-stack of germlines were acquired every 0.25μm, images correspond to a projection using Max intensity method and Fiji macros. Antisera and the dilutions used were as followed: rabbit anti H3K4me3 (Diagenode, 15310003[CS-003100]; 1:12000) and anti-rabbit Alexa Fluor 555 (Invitrogen/Molecular probes #A21428; 1:1000).

### Scoring condensin-I RNAi animals

For scoring the efficacy of RNAi, wildtype or *set-2(bn129)* L4 stage animals were transferred to the same RNAi plates used for scoring germline phenotypes and allowed to develop into adults at 20°C. Following 24hrs of egg laying, animals were removed and the body length of F1 progeny measured 3 days later, at the L4 stage (based on vulval morphology). Animals were placed on 4% agar pads in 1X M9 complemented with 10mM Levamisole, and observed by DIC on an Axio Imager A2 microscope and EC plan Neofluar 10X/0.5 objective. Length of individual worms was measured using Fiji macros.

### Scoring embryonic viability

10-20 L4 hermaphrodites grown at the permissive temperature (15°C), were placed on NGM plates at the restrictive temperature of 20°C during 24h, then removed. Eggs laid were allowed to develop during 24h at 20°C (Normal development takes 18h at 20°C from fertilization to hatching). Then dead eggs and larvae were recovered in 1X M9 and washed twice in 1X M9 in order to remove bacteria. Eggs and larvae were placed on 4% agar pads and a coverslip was placed on top. The stage of embryos was determined by DIC observation with EC Plan-Neofluar 100X/1.3 oil objective and AxioImager A2 (Zeiss). At least 75 embryos were scored per genotypes, and the experiment repeated 3 independent times.

### RNA sequencing of dissected gonads

Gonad dissections and extractions were performed as in (Robert *et al*, 2014). Briefly, prior to dissection worms were placed on NGM plates without food to expel bacteria from the gut. Gonads of 5 to 7 young adults at the L4 stage + 12 h were dissected in dissection buffer (Egg Buffer1.1 X (HEPES pH 7.3 25 mM, NaCl 118 mM, KCL 48 mM, CaCl2 2 mM, 2 mM MgCl2), 0.5 mM Levamisole, 0.1% Tween 20) on slides. Extruded gonads were cut at the elbow and the distal part recovered using a drawn capillary and transferred to 30 μl of XB extraction buffer (Kit Picopure, Life technology, # 12204-01), frozen in liquid nitrogen and stored at −80 ° C. For RNA preparation tubes were thawed, the volume of XB extraction buffer adjusted to 100 μl, and RNA purified using the PicoPure kit (Life Technology, # 12204-01) according to the manufacturer’s instructions. Elution was in 13 μl of nuclease-free water. The integrity of RNA was evaluated using Tape Station 4200 (Agilent), and the concentration of RNA measured using DropSense 96 (Trinean). Construction of rRNA depleted libraries was carried out at the GenomEast platform (IGBMC, Strasbourg, France), and sequencing by an Illumina Hiseq 4000 device.

Bioinformatic analysis was carried out under Galaxy (Afgan *et al*, 2018). Sequence reads were mapped onto the reference genome (WS254) with the RNA-STAR tool (Version 2.4.1d). Sequences with a quality of cartography lower than 10 were removed with SAMtools (Version0.1.19). The expression level of each gene for each sample was calculated with htseq-count (Version 0.7.2). Differential analysis of gene expression between the different strains was carried out with the DESeq2 package version 1.16.1 (Love *et al*, 2014) under R version 3.4.4. Additional analyzes were performed with R.

**Table S1. Genes misregulated in gonads from *set-2* mutant animals**

**Figure S1.**
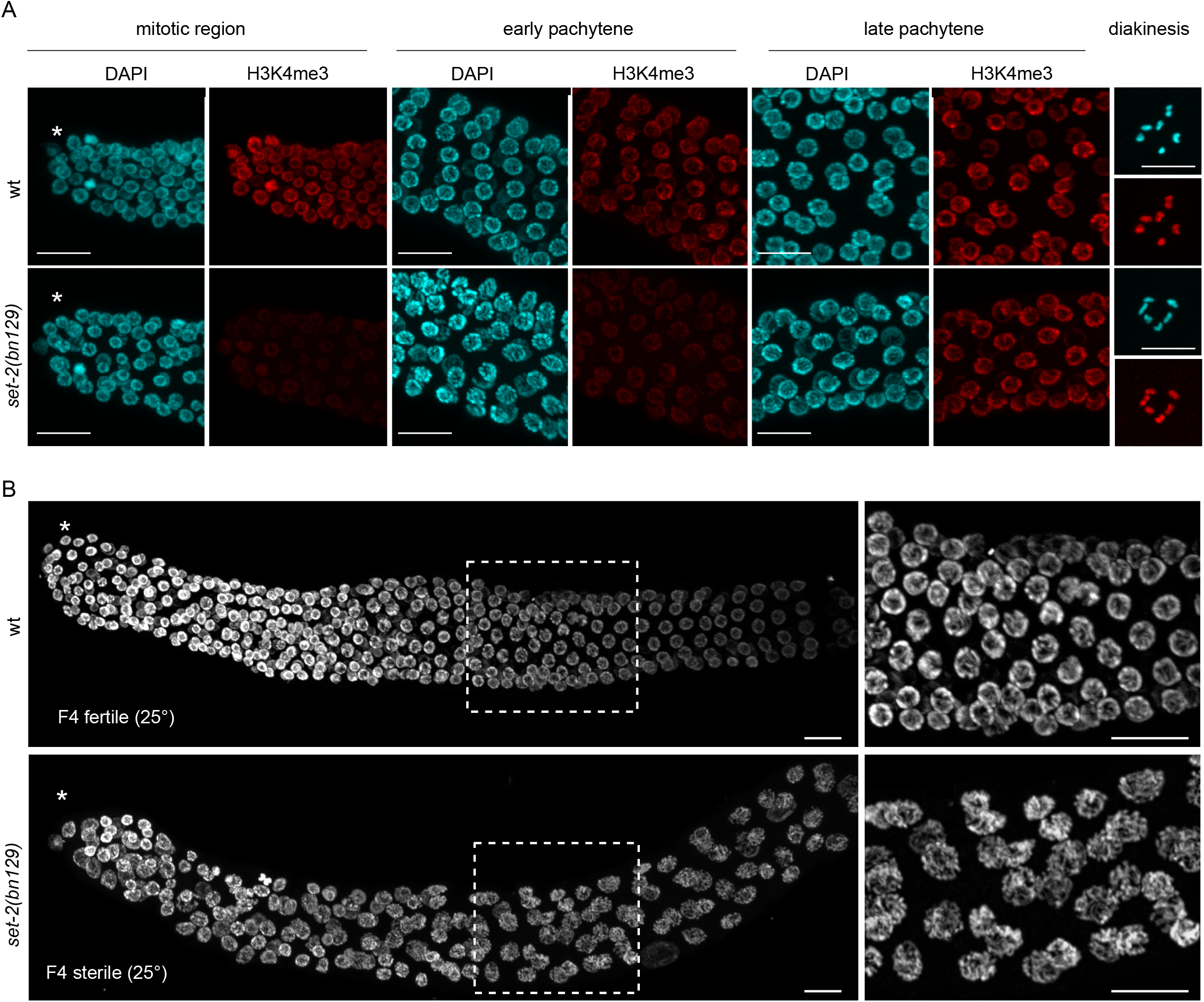
*set-2* inactivation differentially impacts H3K4me3 in the germline. (**A**) Z-projection of confocal images through the mitotic region, early-mid pachytene and late pachytene nuclei, and diakinesis. Gonads were dissected, fixed and probed with rabbit anti-H3K4me3 antibodies (Diagenode, #15310003) and counter-stained with DAPI. Images were taken with the same laser parameter for each condition; scale bar =10 μm. (**B**) Confocal images of DAPI stained germlines of fertile wildtype animals and sterile s*et-2(bn129)* after 4 generations at 25°C; Scale bar, 10 μm). (*) marks distal end of the gonad; scale bar =10 μm

**Figure S2.**
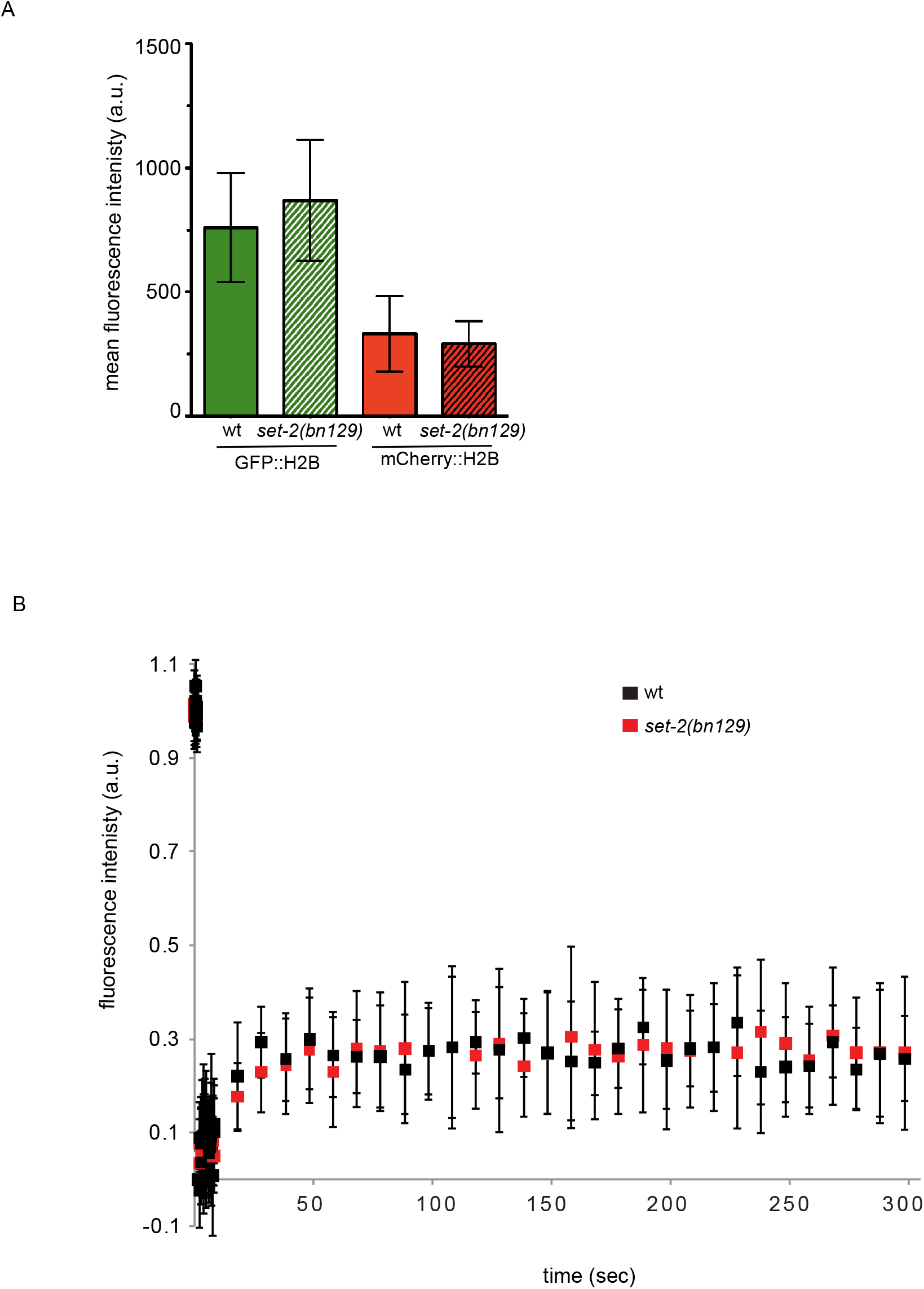
Fluorophore tagged histones H2B are correctly expressed and incorporated into chromatin in *set-2* mutants. (**A**) Mean fluorescence intensity of fluorophore-tagged H2B histones (GFP-H2B and mCherry-H2B) from wt and *set-2(bn129)* mutant pachytene cells. Each data point shows the mean fluorescence intensity ± S.D. from 8 and 12 gonads representing 549 and 698 nuclei from wt and *set-2(bn129)*, respectively. (**B**) Normalized fluorescence intensity recovery of GFP-H2B from wt (black squares) and *set-2(bn129)* (red squares) pachytene cells after laser photo-bleaching experiments (FRAP). Each data point shows the mean ± S.D. for 11 and 8 pachytene cells from wt and *set-2(bn129)*, respectively.

**Figure S3.**
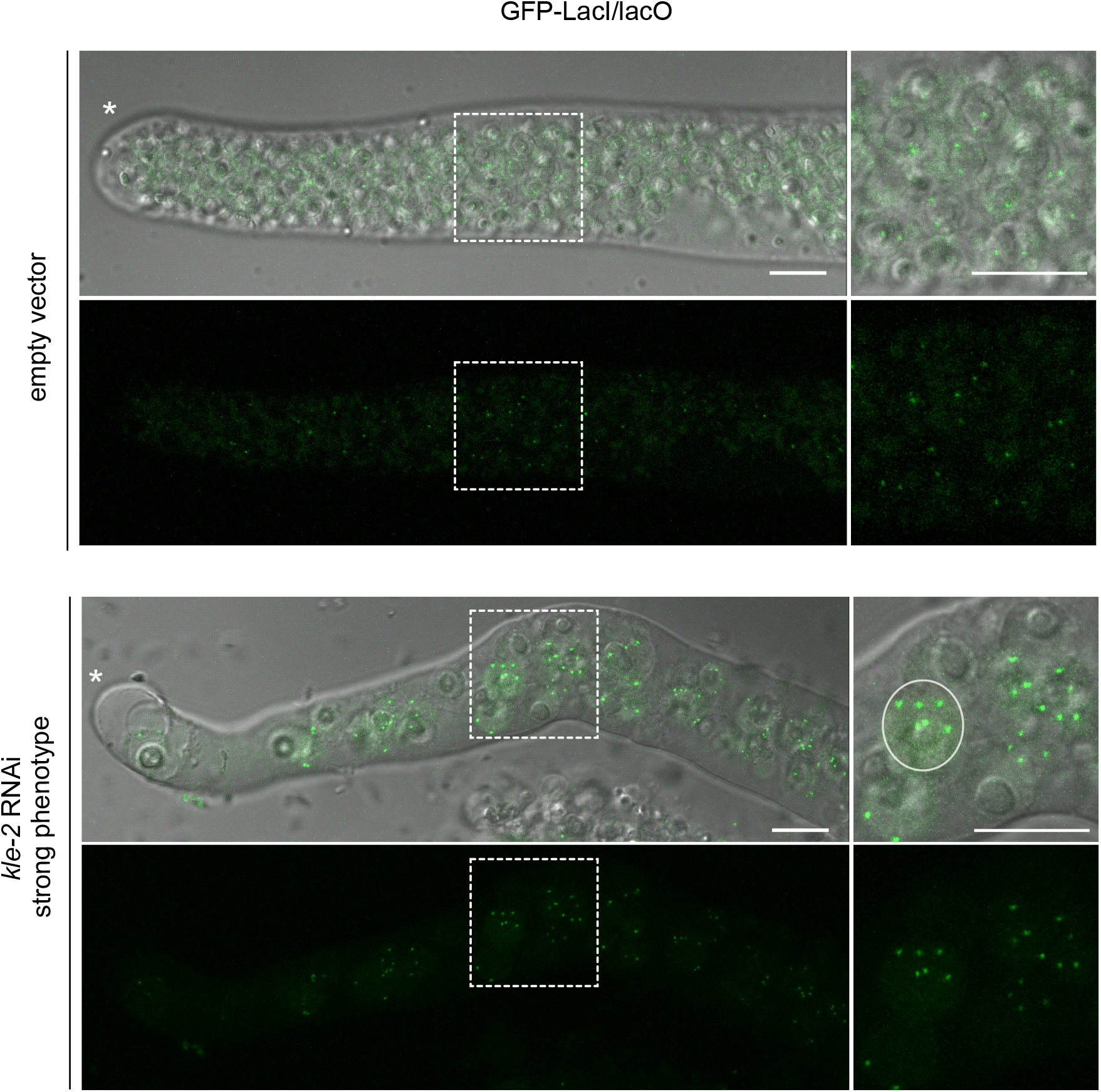
Polyploid cells resulting from *kle-2* knockdown in *set-2* mutant germlines. Wildtype diploid cells carrying a GFP LacO cassette integrated on chromosome IV and expressing lacI show 2 GFP spots/nucleus. Polyploid nuclei from *kle-2*(RNAi);*set-2* germlines showing an “abnormal” phenotype are larger and show more than two GFP spots (circled in white); scale bar=10 μm. (*) marks distal end of the gonad.

**Figure S4.**
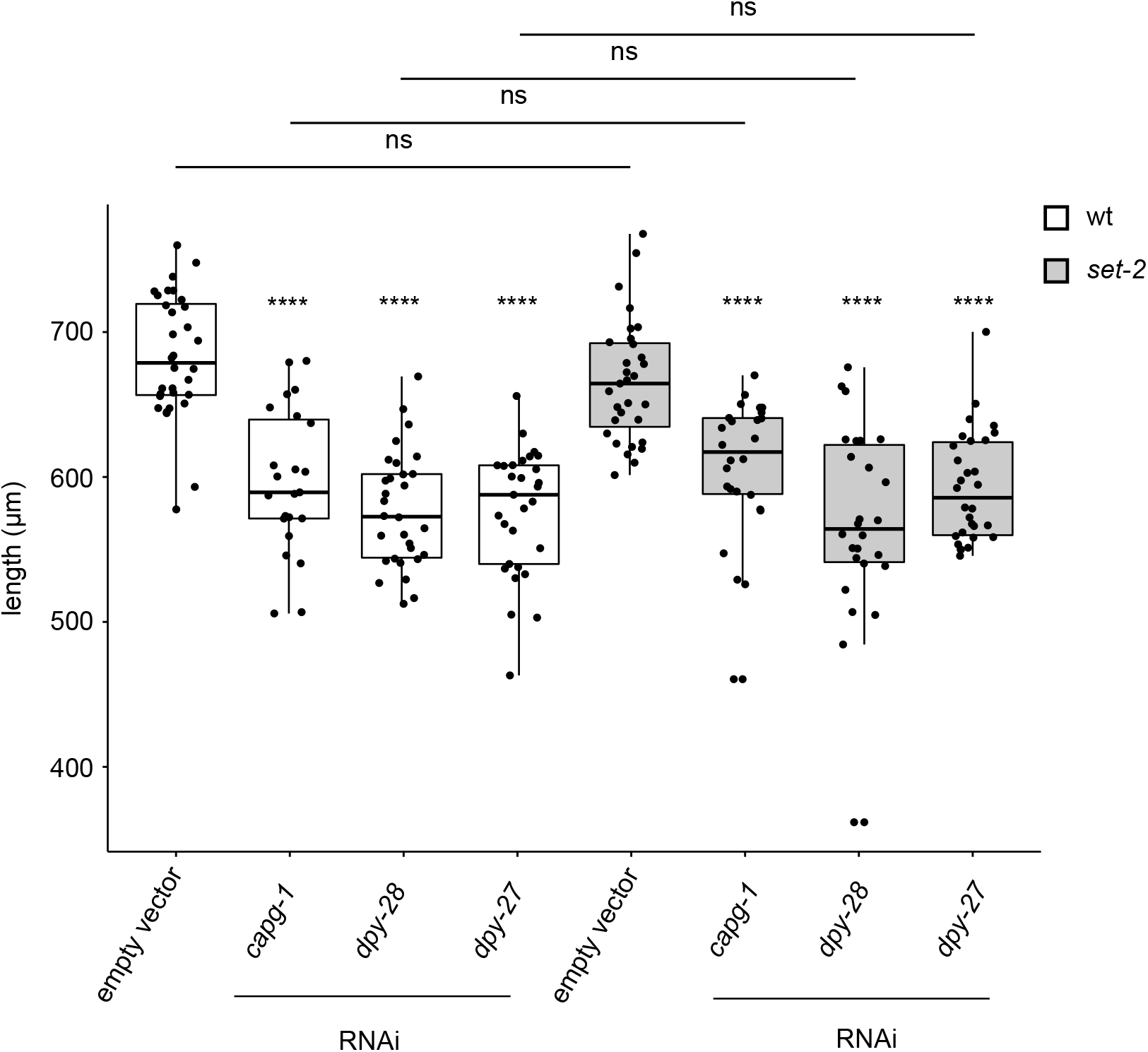
Condensin-I depletion in wildtype and *set-2* mutant animals results in similar reduction in body size. Condensin I knock down animals are smaller in length than control empty vector (dpy phenotype (Zhuang *et al*, 2013). A similar decrease in body size was observed in wildtype and *set-2* mutants. [****] p <0.0001 (t-test). ns, not significant. For wild type: empty vector n=32, *capg-1* RNAi n=23, *dpy-27* RNAi n=29, *dpy-28* RNAi n=30; for *set-2(bn129)*: empty vector n=31, *capg-1* RNAi n=26, *dpy-27* RNAi n=30, *dpy-28* RNAi n=26

**Figure S5.**
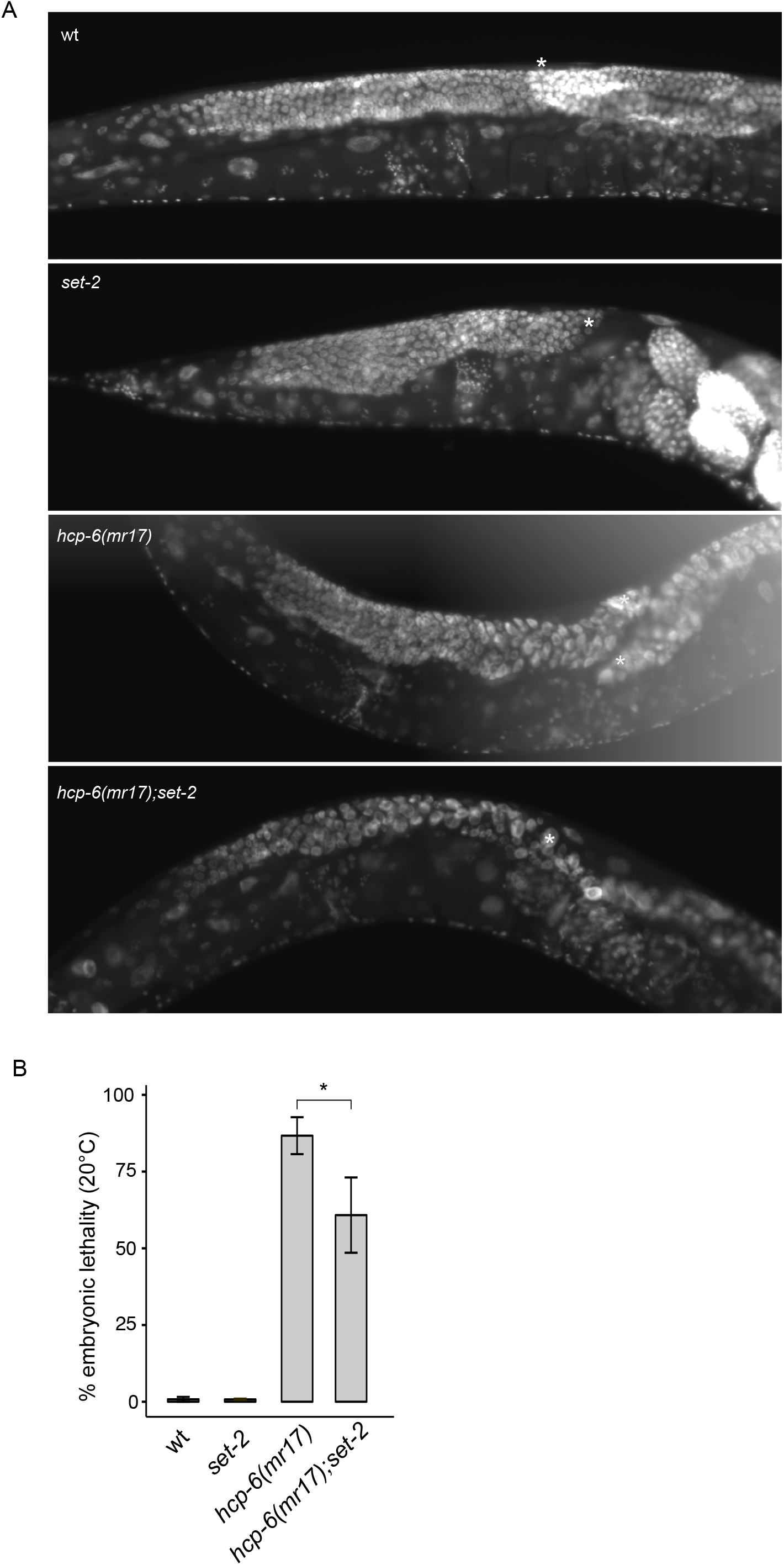
Germline defects of *hcp-6(mr17)* mutant animals and suppression of *hcp-6(mr17)* embryonic lethality. (**A**) *hcp-6(mr17)* gonads are highly disorganized. Young adult wildtype, *set-2*, *hcp-6* and *set-2;hpc-6* mutant animals shifted to 25°C at the L4 stage were DAPI stained and observed on AxioImager A2 (Zeiss). (*) marks distal end of the gonad. (**B**) *set-2* partially suppresses the embryonic lethality of *hcp-6(mr17)* mutants. Percentage of embryonic lethality at 20°C. [*] p < 0.05 (t-test). number of progeny scored for wildtype, *set-2(bn129)*, *hcp-6(mr17)* and *hcp-6(mr17)*;*set-2(bn129)* mutants was 10109, 8227, 8757 and 3961, respectively. 3 independent biological replicates were performed for each genotype.

**Figure S6.**
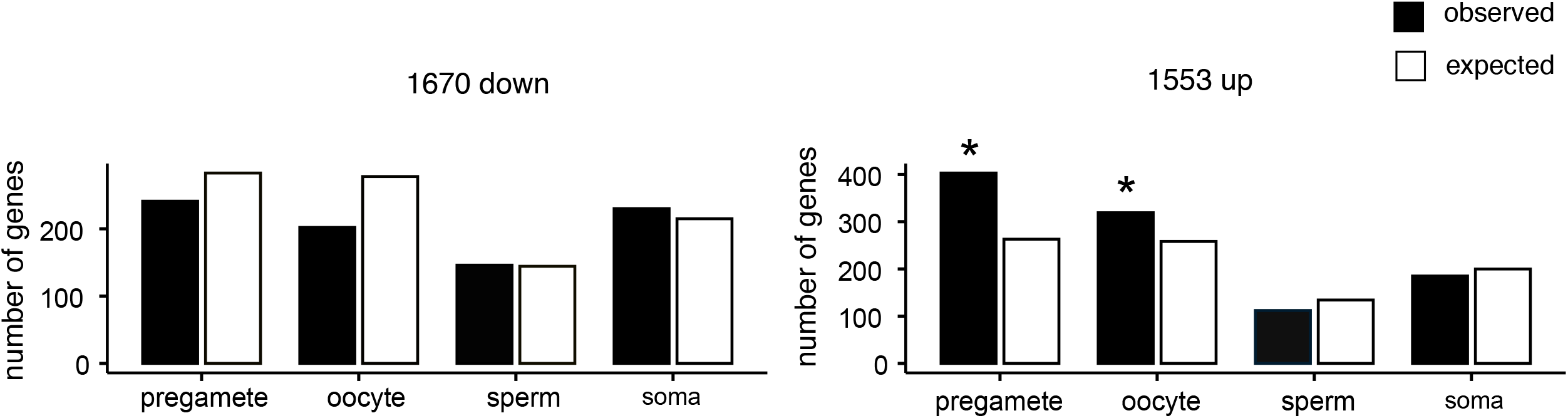
Distribution of *set-2* misregulated genes in different expression classes. Bar graphs indicating expected and observed number of genes in different genes categories based on their preferential expression pattern defined by (Lee *et al*, 2017). Asterisks indicate significantly more genes than expected using an hypergeometric test (p-value <0.001 [*]). Expected values were calculated as: (number of significantly up or down regulated genes) x (number of genes in each category)/number of expressed genes (9377).

**Figure S7.**
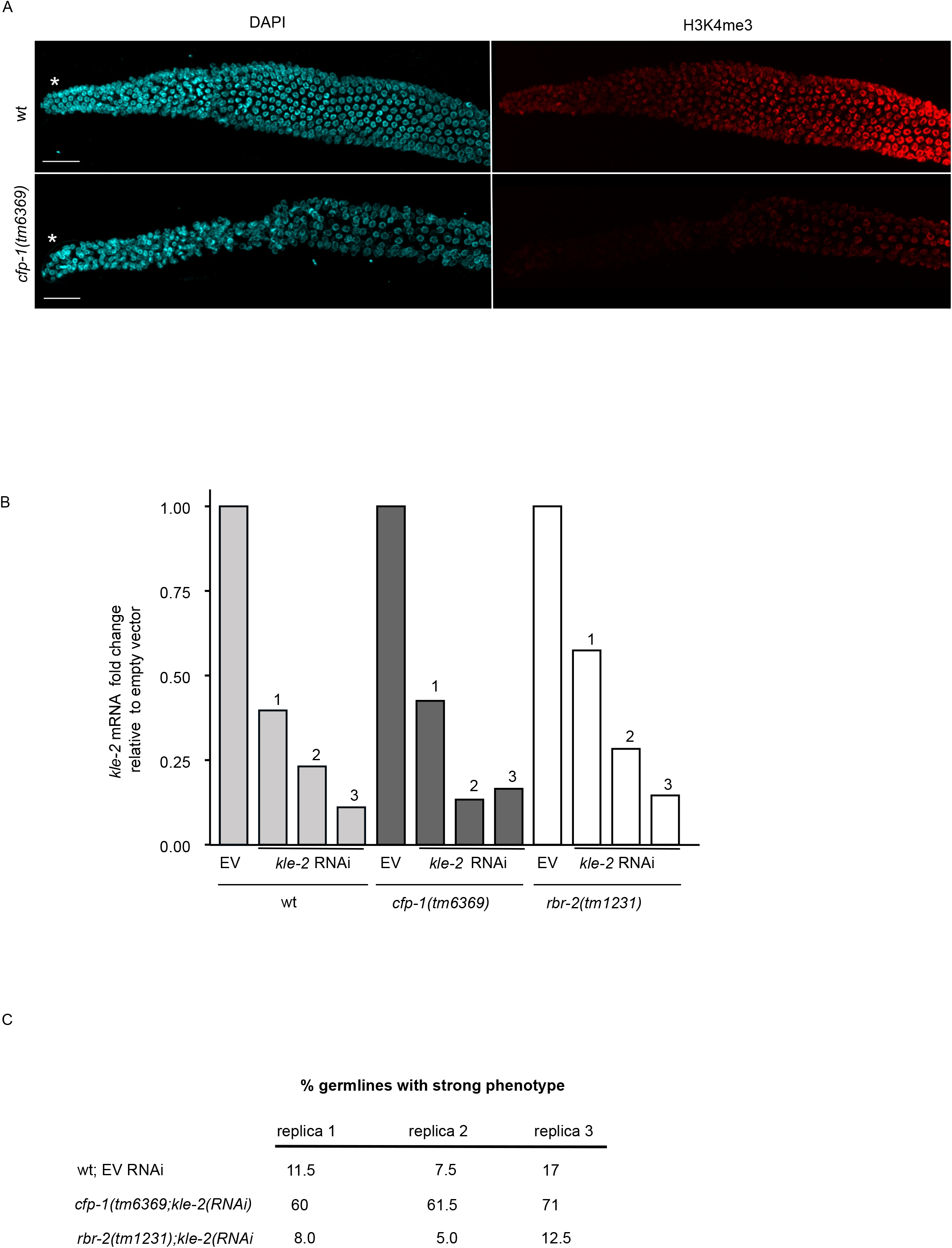
Loss of H3K4me3 in *cfp-1* mutant germline and efficacy of *cfp-1* and *rbr-2* RNAi in independent experiments. (**A**) Decreased H3K4me3 in *cfp-1(tm6369)* mutant germline. (*) marks distal end of the gonad. (**B**) *kle-2* mRNA levels in wt, *cfp-1(tm6369)* and *rbr-2(tm1231)* mutant animals. For each of the three independent experiments, relative fold change was calculated with respect to empty vector condition, following normalization with *pmp-3* and *cdc-42*. (**C**) Percentage of germlines with “abnormal” phenotype in each of the three replicas after RNAi directed against *kle-2* RNAi. Note that although the efficacy of *kle-2* mRNA knockdown varied between the three experiments, the percentage of abnormal germlines scored for each genotype (wt, *cfp-1* and *rbr-2*) was highly similar.

## References

Acquaviva L, Szekvolgyi L, Dichtl B, Dichtl BS, de La Roche Saint Andre C, Nicolas A & Geli V (2013) The COMPASS subunit Spp1 links histone methylation to initiation of meiotic recombination. Science 339: 215–8

Adachi Y, Luke M & Laemmli UK (1991) Chromosome assembly in vitro: topoisomerase II is required for condensation. Cell 64: 137–148

Adriaens C, Serebryannyy LA, Feric M, Schibler A, Meaburn KJ, Kubben N, Trzaskoma P, Shachar S, Vidak S, Finn EH, Sood V, Pegoraro G & Misteli T (2018) Blank spots on the map: some current questions on nuclear organization and genome architecture. Histochem. Cell Biol. 150: 579–592

Afgan E, Baker D, Batut B, van den Beek M, Bouvier D, Cech M, Chilton J, Clements D, Coraor N, Grüning BA, Guerler A, Hillman-Jackson J, Hiltemann S, Jalili V, Rasche H, Soranzo N, Goecks J, Taylor J, Nekrutenko A & Blankenberg D (2018) The Galaxy platform for accessible, reproducible and collaborative biomedical analyses: 2018 update. Nucleic Acids Res. 46: W537–W544

Alavattam KG, Maezawa S, Sakashita A, Khoury H, Barski A, Kaplan N & Namekawa SH (2019) Attenuated chromatin compartmentalization in meiosis and its maturation in sperm development. Nat. Struct. Mol. Biol. 26: 175–184

Albritton SE & Ercan S (2018) Caenorhabditis elegans Dosage Compensation: Insights into Condensin-Mediated Gene Regulation. Trends Genet. TIG 34: 41–53

Alvares SM, Mayberry GA, Joyner EY, Lakowski B & Ahmed S (2014) H3K4 demethylase activities repress proliferative and postmitotic aging. Aging Cell 13: 245–253

Antonin W & Neumann H (2016) Chromosome condensation and decondensation during mitosis. Curr. Opin. Cell Biol. 40: 15–22

Baarlink C, Plessner M, Sherrard A, Morita K, Misu S, Virant D, Kleinschnitz E-M, Harniman R, Alibhai D, Baumeister S, Miyamoto K, Endesfelder U, Kaidi A & Grosse R (2017) A transient pool of nuclear F-actin at mitotic exit controls chromatin organization. Nat. Cell Biol. 19: 1389–1399

Beck DB, Oda H, Shen SS & Reinberg D (2012) PR-Set7 and H4K20me1: at the crossroads of genome integrity, cell cycle, chromosome condensation, and transcription. Genes Dev. 26: 325–337

Beurton F, Stempor P, Caron M, Appert A, Dong Y, Chen RA-J, Cluet D, Couté Y, Herbette M, Huang N, Polveche H, Spichty M, Bedet C, Ahringer J & Palladino F (2019) Physical and functional interaction between SET1/COMPASS complex component CFP-1 and a Sin3S HDAC complex in C. elegans. Nucleic Acids Res.

Blat Y, Protacio RU, Hunter N & Kleckner N (2002) Physical and functional interactions among basic chromosome organizational features govern early steps of meiotic chiasma formation. Cell 111: 791–802

Borde V, Robine N, Lin W, Bonfils S, Geli V & Nicolas A (2009) Histone H3 lysine 4 trimethylation marks meiotic recombination initiation sites. EMBO J 28: 99–111

Boulton SJ, Martin JS, Polanowska J, Hill DE, Gartner A & Vidal M (2004) BRCA1/BARD1 orthologs required for DNA repair in Caenorhabditis elegans. Curr Biol 14: 33–9

Brenner S (1974) The Genetics of Caenorhabditis elegans. Genetics 77: 71–94

Buard J, Barthès P, Grey C & de Massy B (2009) Distinct histone modifications define initiation and repair of meiotic recombination in the mouse. EMBO J. 28: 2616–2624

Buchenau P, Saumweber H & Arndt-Jovin DJ (1993) Consequences of topoisomerase II inhibition in early embryogenesis of Drosophila revealed by in vivo confocal laser scanning microscopy. J. Cell Sci. 104 (Pt 4): 1175–1185

Chan RC, Severson AF & Meyer BJ (2004) Condensin restructures chromosomes in preparation for meiotic divisions. J. Cell Biol. 167: 613–625

Chen RA-J, Stempor P, Down TA, Zeiser E, Feuer SK & Ahringer J (2014) Extreme HOT regions are CpG-dense promoters in C. elegans and humans. Genome Res. 24: 1138–1146

Christensen J, Agger K, Cloos PA, Pasini D, Rose S, Sennels L, Rappsilber J, Hansen KH, Salcini AE & Helin K (2007) RBP2 belongs to a family of demethylases, specific for tri-and dimethylated lysine 4 on histone 3. Cell 128: 1063–76

Clouaire T, Webb S & Bird A (2014) Cfp1 is required for gene expression-dependent H3K4 trimethylation and H3K9 acetylation in embryonic stem cells. Genome Biol. 15: 451

Clouaire T, Webb S, Skene P, Illingworth R, Kerr A, Andrews R, Lee J-H, Skalnik D & Bird A (2012) Cfp1 integrates both CpG content and gene activity for accurate H3K4me3 deposition in embryonic stem cells. Genes Dev. 26: 1714–1728

Csankovszki G, Collette K, Spahl K, Carey J, Snyder M, Petty E, Patel U, Tabuchi T, Liu H, McLeod I, Thompson J, Sarkesik A, Yates J, Meyer BJ & Hagstrom K (2009) Three Distinct Condensin Complexes Control C. elegans Chromosome Dynamics. Curr. Biol. 19: 9–19

Cuvier O & Hirano T (2003) A role of topoisomerase II in linking DNA replication to chromosome condensation. J. Cell Biol. 160: 645–655

Dehe PM, Dichtl B, Schaft D, Roguev A, Pamblanco M, Lebrun R, Rodriguez-Gil A, Mkandawire M, Landsberg K, Shevchenko A, Shevchenko A, Rosaleny LE, Tordera V, Chavez S, Stewart AF & Geli V (2006) Protein interactions within the Set1 complex and their roles in the regulation of histone 3 lysine 4 methylation. J Biol Chem 281: 35404–12

DiNardo S, Voelkel K & Sternglanz R (1984) DNA topoisomerase II mutant of Saccharomyces cerevisiae: topoisomerase II is required for segregation of daughter molecules at the termination of DNA replication. Proc. Natl. Acad. Sci. 81: 2616–2620

Eissenberg JC & Shilatifard A (2010) Histone H3 lysine 4 (H3K4) methylation in development and differentiation. Dev Biol 339: 240–9

Fisher K, Southall SM, Wilson JR & Poulin GB Methylation and demethylation activities of a C. elegans MLL-like complex attenuate RAS signalling. Dev Biol 341: 142–53

Ganji M, Shaltiel IA, Bisht S, Kim E, Kalichava A, Haering CH & Dekker C (2018) Real-time imaging of DNA loop extrusion by condensin. Science 360: 102–105

Gibcus JH, Samejima K, Goloborodko A, Samejima I, Naumova N, Nuebler J, Kanemaki MT, Xie L, Paulson JR, Earnshaw WC, Mirny LA & Dekker J (2018) A pathway for mitotic chromosome formation. Science 359:

Gómez R, Viera A, Berenguer I, Llano E, Pendás AM, Barbero JL, Kikuchi A & Suja JA (2014) Cohesin removal precedes topoisomerase IIα-dependent decatenation at centromeres in male mammalian meiosis II. Chromosoma 123: 129–146

Gonzalez-Serricchio AS & Sternberg PW (2006) Visualization of C. elegans transgenic arrays by GFP. BMC Genet. 7: 36

Gumienny TL, Lambie E, Hartwieg E, Horvitz HR & Hengartner MO (1999) Genetic control of programmed cell death in the Caenorhabditis elegans hermaphrodite germline. Development 126: 1011–22

Heestand B, Simon M, Frenk S, Titov D & Ahmed S (2018) Transgenerational Sterility of Piwi Mutants Represents a Dynamic Form of Adult Reproductive Diapause. Cell Rep. 23: 156–171

Herbette M, Mercier MG, Michal F, Cluet D, Burny C, Yvert G, Robert VJ & Palladino F (2017) The C. elegans SET-2/SET1 histone H3 Lys4 (H3K4) methyltransferase preserves genome stability in the germline. DNA Repair 57: 139–150

Hirano T (2012) Condensins: universal organizers of chromosomes with diverse functions. Genes Dev. 26: 1659–1678

Hirano T & Mitchison TJ (1993) Topoisomerase II does not play a scaffolding role in the organization of mitotic chromosomes assembled in Xenopus egg extracts. J. Cell Biol. 120: 601–612

Hong Y, Sonneville R, Agostinho A, Meier B, Wang B, Blow JJ & Gartner A (2016) The SMC-5/6 Complex and the HIM-6 (BLM) Helicase Synergistically Promote Meiotic Recombination Intermediate Processing and Chromosome Maturation during Caenorhabditis elegans Meiosis. PLoS Genet. 12: e1005872

Howe FS, Fischl H, Murray SC & Mellor J (2017) Is H3K4me3 instructive for transcription activation? BioEssays News Rev. Mol. Cell. Dev. Biol. 39: 1–12

Hudson DF, Vagnarelli P, Gassmann R & Earnshaw WC (2003) Condensin Is Required for Nonhistone Protein Assembly and Structural Integrity of Vertebrate Mitotic Chromosomes. Dev. Cell 5: 323–336

Hughes SE & Hawley RS (2014) Topoisomerase II is required for the proper separation of heterochromatic regions during Drosophila melanogaster female meiosis. PLoS Genet. 10: e1004650

Jabbari K, Wirtz J, Rauscher M & Wiehe T (2019) A common genomic code for chromatin architecture and recombination landscape. PloS One 14: e0213278

Jaramillo-Lambert A, Fabritius AS, Hansen TJ, Smith HE & Golden A (2016) The Identification of a Novel Mutant Allele of topoisomerase II in Caenorhabditis elegans Reveals a Unique Role in Chromosome Segregation During Spermatogenesis. Genetics 204: 1407–1422

Kadyk LC & Kimble J (1998) Genetic regulation of entry into meiosis in Caenorhabditis elegans. Dev. Camb. Engl. 125: 1803–1813

Kim H, Yen L, Wongpalee SP, Kirshner JA, Mehta N, Xue Y, Johnston JB, Burlingame AL, Kim JK, Loparo JJ & Jacobsen SE (2019) The Gene-Silencing Protein MORC-1 Topologically Entraps DNA and Forms Multimeric Assemblies to Cause DNA Compaction. Mol. Cell 75: 700–710.e6

Kimble J & Crittenden SL (2005) Germline proliferation and its control. WormBook Online Rev. C Elegans Biol.: 1–14

Klein F (1992) Localization of RAP1 and topoisomerase II in nuclei and meiotic chromosomes of yeast. J. Cell Biol. 117: 935–948

Kruitwagen T, Denoth-Lippuner A, Wilkins BJ, Neumann H & Barral Y (2015) Axial contraction and short-range compaction of chromatin synergistically promote mitotic chromosome condensation. eLife 4: e1039

Ladouceur A-M, Ranjan R, Smith L, Fadero T, Heppert J, Goldstein B, Maddox AS & Maddox PS (2017) CENP-A and topoisomerase-II antagonistically affect chromosome length. J. Cell Biol. 216: 2645–2655

Lam K-WG, Brick K, Cheng G, Pratto F & Camerini-Otero RD (2019) Cell-type-specific genomics reveals histone modification dynamics in mammalian meiosis. Nat. Commun. 10: 3821

Lee C-YS, Lu T & Seydoux G (2017) Nanos promotes epigenetic reprograming of the germline by down-regulation of the THAP transcription factor LIN-15B. eLife 6:

Lenstra TL, Benschop JJ, Kim T, Schulze JM, Brabers NA, Margaritis T, van de Pasch LA, van Heesch SA, Brok MO, Groot Koerkamp MJ, Ko CW, van Leenen D, Sameith K, van Hooff SR, Lijnzaad P, Kemmeren P, Hentrich T, Kobor MS, Buratowski S & Holstege FC (2011) The specificity and topology of chromatin interaction pathways in yeast. Mol Cell 42: 536–49

Li T & Kelly WG (2011) A role for Set1/MLL-related components in epigenetic regulation of the Caenorhabditis elegans germ line. PLoS Genet 7: e1001349

Li X-M, Yu C, Wang Z-W, Zhang Y-L, Liu X-M, Zhou D, Sun Q-Y & Fan H-Y (2013) DNA Topoisomerase II Is Dispensable for Oocyte Meiotic Resumption but Is Essential for Meiotic Chromosome Condensation and Separation in Mice1. Biol. Reprod. 89: Available at: https://academic.oup.com/biolreprod/article-lookup/doi/10.1095/biolreprod.113.110692 [Accessed August 22, 2019]

Liang Z, Zickler D, Prentiss M, Chang FS, Witz G, Maeshima K & Kleckner N (2015) Chromosomes Progress to Metaphase in Multiple Discrete Steps via Global Compaction/Expansion Cycles. Cell 161: 1124–1137

Llères D, Bailly AP, Perrin A, Norman DG, Xirodimas DP & Feil R (2017) Quantitative FLIM-FRET Microscopy to Monitor Nanoscale Chromatin Compaction In Vivo Reveals Structural Roles of Condensin Complexes. Cell Rep. 18: 1791–1803

Llères D, James J, Swift S, Norman DG & Lamond AI (2009) Quantitative analysis of chromatin compaction in living cells using FLIM–FRET. J. Cell Biol. 187: 481–496

Llères D, Swift S & Lamond AI (2007) Detecting Protein-Protein Interactions In Vivo with FRET using Multiphoton Fluorescence Lifetime Imaging Microscopy (FLIM). Curr. Protoc. Cytom. 42: Available at: https://onlinelibrary.wiley.com/doi/abs/10.1002/0471142956.cy1210s42 [Accessed February 28, 2020]

Lorenz DR, Mikheyeva IV, Johansen P, Meyer L, Berg A, Grewal SIS & Cam HP (2012) CENP-B cooperates with Set1 in bidirectional transcriptional silencing and genome organization of retrotransposons. Mol. Cell. Biol. 32: 4215–4225

Lou J, Scipioni L, Wright BK, Bartolec TK, Zhang J, Masamsetti VP, Gaus K, Gratton E, Cesare AJ & Hinde E (2019) Phasor histone FLIM-FRET microscopy quantifies spatiotemporal rearrangement of chromatin architecture during the DNA damage response. Proc. Natl. Acad. Sci. 116: 7323–7332

Love MI, Huber W & Anders S (2014) Moderated estimation of fold change and dispersion for RNA-seq data with DESeq2. Genome Biol. 15: 550

MacGregor IA, Adams IR & Gilbert N (2019) Large-scale chromatin organisation in interphase, mitosis and meiosis. Biochem. J. 476: 2141–2156

Marchetti F, Bishop JB, Lowe X, Generoso WM, Hozier J & Wyrobek AJ (2001) Etoposide induces heritable chromosomal aberrations and aneuploidy during male meiosis in the mouse. Proc. Natl. Acad. Sci. 98: 3952–3957

Markaki Y, Christogianni A, Politou AS & Georgatos SD (2009) Phosphorylation of histone H3 at Thr3 is part of a combinatorial pattern that marks and configures mitotic chromatin. J. Cell Sci. 122: 2809–2819

McNicoll F, Stevense M & Jessberger R (2013) Cohesin in Gametogenesis. In Current Topics in Developmental Biology pp 1–34. Elsevier Available at: https://linkinghub.elsevier.com/retrieve/pii/B9780124160248000015 [Accessed October 9, 2019]

Mets DG & Meyer BJ (2009) Condensins regulate meiotic DNA break distribution, thus crossover frequency, by controlling chromosome structure. Cell 139: 73–86

Mikheyeva IV, Grady PJR, Tamburini FB, Lorenz DR & Cam HP (2014) Multifaceted genome control by Set1 Dependent and Independent of H3K4 methylation and the Set1C/COMPASS complex. PLoS Genet. 10: e1004740

Nagasaka K, Hossain MJ, Roberti MJ, Ellenberg J & Hirota T (2016) Sister chromatid resolution is an intrinsic part of chromosome organization in prophase. Nat. Cell Biol. 18: 692–699

O’Connell KF, Leys CM & White JG (1998) A genetic screen for temperature-sensitive celldivision mutants of Caenorhabditis elegans. Genetics 149: 1303–1321

Oliveira RA, Hamilton RS, Pauli A, Davis I & Nasmyth K (2010) Cohesin cleavage and Cdk inhibition trigger formation of daughter nuclei. Nat. Cell Biol. 12: 185–192

Ono T, Fang Y, Spector DL & Hirano T (2004) Spatial and temporal regulation of Condensins I and II in mitotic chromosome assembly in human cells. Mol. Biol. Cell 15: 3296–3308

Panizza S, Mendoza MA, Berlinger M, Huang L, Nicolas A, Shirahige K & Klein F (2011) Spo11-accessory proteins link double-strand break sites to the chromosome axis in early meiotic recombination. Cell 146: 372–383

Papaluca A & Ramotar D (2016) A novel approach using C. elegans DNA damage-induced apoptosis to characterize the dynamics of uptake transporters for therapeutic drug discoveries. Sci. Rep. 6: 36026

Patel L, Kang R, Rosenberg SC, Qiu Y, Raviram R, Chee S, Hu R, Ren B, Cole F & Corbett KD (2019) Dynamic reorganization of the genome shapes the recombination landscape in meiotic prophase. Nat. Struct. Mol. Biol. 26: 164–174

Petrova B, Dehler S, Kruitwagen T, Hériché J-K, Miura K & Haering CH (2013) Quantitative analysis of chromosome condensation in fission yeast. Mol. Cell. Biol. 33: 984–998

Piskadlo E & Oliveira RA (2016) Novel insights into mitotic chromosome condensation. F1000Research 5:

Potapova T & Gorbsky GJ (2017) The Consequences of Chromosome Segregation Errors in Mitosis and Meiosis. Biology 6:

Prakash K, Fournier D, Redl S, Best G, Borsos M, Tiwari VK, Tachibana-Konwalski K, Ketting RF, Parekh SH, Cremer C & Birk UJ (2015) Superresolution imaging reveals structurally distinct periodic patterns of chromatin along pachytene chromosomes. Proc. Natl. Acad. Sci. 112: 14635–14640

Robert VJ, Mercier MG, Bedet C, Janczarski S, Merlet J, Garvis S, Ciosk R & Palladino F (2014) The SET-2/SET1 histone H3K4 methyltransferase maintains pluripotency in the Caenorhabditis elegans germline. Cell Rep 9: 443–50

Rowley MJ & Corces VG (2018) Organizational principles of 3D genome architecture. Nat. Rev. Genet. 19: 789–800

Rowley MJ, Nichols MH, Lyu X, Ando-Kuri M, Rivera ISM, Hermetz K, Wang P, Ruan Y & Corces VG (2017) Evolutionarily Conserved Principles Predict 3D Chromatin Organization. Mol. Cell 67: 837–852.e7

Sakaguchi A & Kikuchi A (2004) Functional compatibility between isoform alpha and beta of type II DNA topoisomerase. J. Cell Sci. 117: 1047–1054

Samejima K, Samejima I, Vagnarelli P, Ogawa H, Vargiu G, Kelly DA, de Lima Alves F, Kerr A, Green LC, Hudson DF, Ohta S, Cooke CA, Farr CJ, Rappsilber J & Earnshaw WC (2012) Mitotic chromosomes are compacted laterally by KIF4 and condensin and axially by topoisomerase IIα. J. Cell Biol. 199: 755–770

Schalbetter SA, Fudenberg G, Baxter J, Pollard KS & Neale MJ (2019) Principles of meiotic chromosome assembly revealed in S. cerevisiae. Nat. Commun. 10: 4795

Schindelin J, Arganda-Carreras I, Frise E, Kaynig V, Longair M, Pietzsch T, Preibisch S, Rueden C, Saalfeld S, Schmid B, Tinevez J-Y, White DJ, Hartenstein V, Eliceiri K, Tomancak P & Cardona A (2012) Fiji: an open-source platform for biological-image analysis. Nat. Methods 9: 676–682

Sha Q-Q, Dai X-X, Jiang J-C, Yu C, Jiang Y, Liu J, Ou X-H, Zhang S-Y & Fan H-Y (2018) CFP1 coordinates histone H3 lysine-4 trimethylation and meiotic cell cycle progression in mouse oocytes. Nat. Commun. 9: 3477

Shintomi K, Takahashi TS & Hirano T (2015) Reconstitution of mitotic chromatids with a minimum set of purified factors. Nat. Cell Biol. 17: 1014–1023

Simonet T, Dulermo R, Schott S & Palladino F (2007) Antagonistic functions of SET-2/SET1 and HPL/HP1 proteins in C. elegans development. Dev Biol 312: 367–83

Sobecki M, Mrouj K, Camasses A, Parisis N, Nicolas E, Llères D, Gerbe F, Prieto S, Krasinska L, David A, Eguren M, Birling M-C, Urbach S, Hem S, Déjardin J, Malumbres M, Jay P, Dulic V, Lafontaine DL, Feil R, et al (2016) The cell proliferation antigen Ki-67 organises heterochromatin. eLife 5: e13722

Sommermeyer V, Beneut C, Chaplais E, Serrentino ME & Borde V (2013) Spp1, a member of the Set1 Complex, promotes meiotic DSB formation in promoters by tethering histone H3K4 methylation sites to chromosome axes. Mol Cell 49: 43–54

Stear JH & Roth MB (2002) Characterization of HCP-6, a C. elegans protein required to prevent chromosome twisting and merotelic attachment. Genes Dev. 16: 1498–1508

T. & Förster (1949) Experimental and theoretical investigation of the intermolecular transfer of electronic excitation energy. Z. Naturforsch. A. 4: 321–327

Uemura T, Ohkura H, Adachi Y, Morino K, Shiozaki K & Yanagida M (1987) DNA topoisomerase II is required for condensation and separation of mitotic chromosomes in S. pombe. Cell 50: 917–925

Vagnarelli P, Hudson DF, Ribeiro SA, Trinkle-Mulcahy L, Spence JM, Lai F, Farr CJ, Lamond AI & Earnshaw WC (2006) Condensin and Repo-Man-PP1 co-operate in the regulation of chromosome architecture during mitosis. Nat. Cell Biol. 8: 1133–1142

Wang Y, Sherrard A, Zhao B, Melak M, Trautwein J, Kleinschnitz E-M, Tsopoulidis N, Fackler OT, Schwan C & Grosse R (2019a) GPCR-induced calcium transients trigger nuclear actin assembly for chromatin dynamics. Nat. Commun. 10: 5271

Wang Y, Wang H, Zhang Y, Du Z, Si W, Fan S, Qin D, Wang M, Duan Y, Li L, Jiao Y, Li Y, Wang Q, Shi Q, Wu X & Xie W (2019b) Reprogramming of Meiotic Chromatin Architecture during Spermatogenesis. Mol. Cell 73: 547–561.e6

Weiner A, Chen HV, Liu CL, Rahat A, Klien A, Soares L, Gudipati M, Pfeffner J, Regev A, Buratowski S, Pleiss JA, Friedman N & Rando OJ (2012) Systematic dissection of roles for chromatin regulators in a yeast stress response. PLoS Biol 10: e1001369

Weiser NE, Yang DX, Feng S, Kalinava N, Brown KC, Khanikar J, Freeberg MA, Snyder MJ, Csankovszki G, Chan RC, Gu SG, Montgomery TA, Jacobsen SE & Kim JK (2017) MORC-1 Integrates Nuclear RNAi and Transgenerational Chromatin Architecture to Promote Germline Immortality. Dev. Cell 41: 408–423.e7

West AM, Rosenberg SC, Ur SN, Lehmer MK, Ye Q, Hagemann G, Caballero I, Usón I, MacQueen AJ, Herzog F & Corbett KD (2019) A conserved filamentous assembly underlies the structure of the meiotic chromosome axis. eLife 8:

Wilkins BJ, Rall NA, Ostwal Y, Kruitwagen T, Hiragami-Hamada K, Winkler M, Barral Y, Fischle W & Neumann H (2014) A Cascade of Histone Modifications Induces Chromatin Condensation in Mitosis. Science 343: 77–80

Xiao Y, Bedet C, Robert VJ, Simonet T, Dunkelbarger S, Rakotomalala C, Soete G, Korswagen HC, Strome S & Palladino F (2011) Caenorhabditis elegans chromatin-associated proteins SET-2 and ASH-2 are differentially required for histone H3 Lys 4 methylation in embryos and adult germ cells. Proc Natl Acad Sci U A 108: 8305–10

Yoshimura SH & Hirano T (2016) HEAT repeats - versatile arrays of amphiphilic helices working in crowded environments? J. Cell Sci. 129: 3963–3970

Yu C, Fan X, Sha Q-Q, Wang H-H, Li B-T, Dai X-X, Shen L, Liu J, Wang L, Liu K, Tang F & Fan H-Y (2017) CFP1 Regulates Histone H3K4 Trimethylation and Developmental Potential in Mouse Oocytes. Cell Rep. 20: 1161–1172

Yuen KC, Slaughter BD & Gerton JL (2017) Condensin II is anchored by TFIIIC and H3K4me3 in the mammalian genome and supports the expression of active dense gene clusters. Sci. Adv. 3: e1700191

Zhiteneva A, Bonfiglio JJ, Makarov A, Colby T, Vagnarelli P, Schirmer EC, Matic I & Earnshaw WC (2017) Mitotic post-translational modifications of histones promote chromatin compaction in vitro. Open Biol. 7:

Zhuang JJ, Banse SA & Hunter CP (2013) The nuclear argonaute NRDE-3 contributes to transitive RNAi in Caenorhabditis elegans. Genetics 194: 117–131

